# Pseudouridine guides germline small RNA transport and epigenetic inheritance

**DOI:** 10.1101/2023.05.27.542553

**Authors:** Rowan P Herridge, Jakub Dolata, Valentina Migliori, Cristiane de Santis Alves, Filipe Borges, Andrea J Schorn, Frédéric Van Ex, Jean-Sebastien Parent, Ann Lin, Mateusz Bajczyk, Tommaso Leonardi, Alan Hendrick, Tony Kouzarides, Robert A Martienssen

**Affiliations:** Howard Hughes Medical Institute, Cold Spring Harbor Laboratory, Cold Spring Harbor, NY, 11724, USA; The Gurdon Institute, University of Cambridge, Tennis Court Road, Cambridge CB2 1QN, UK; Wellcome Sanger Institute, Wellcome Genome Campus, Hinxton, Cambridge, UK; Department of Gene Expression, Institute of Molecular Biology and Biotechnology, Adam Mickiewicz University, Poznan, Poland; Center for Genomic Science of IIT@SEMM, Instituto Italiano di Tecnologia (IIT), 20139 Milan, Italy; Storm Therapeutics, Ltd., Moneta Building (B280), Babraham Research Campus, Cambridge CB22 3AT, UK

## Abstract

Epigenetic modifications that arise during plant and animal development, such as DNA and histone modification, are mostly reset during gamete formation, but some are inherited from the germline including those marking imprinted genes^1^. Small RNAs guide these epigenetic modifications, and some are also inherited by the next generation^2,3^. In *C. elegans*, these inherited small RNAs have poly (UG) tails^4^, but how inherited small RNAs are distinguished in other animals and plants is unknown. Pseudouridine (Ψ) is the most abundant RNA modification but has not been explored in small RNAs. Here, we develop novel assays to detect Ψ in short RNA sequences, demonstrating its presence in mouse and *Arabidopsis* microRNAs and their precursors. We also detect substantial enrichment in germline small RNAs, namely epigenetically activated siRNAs (easiRNAs) in *Arabidopsis* pollen, and piwi-interacting piRNAs in mouse testis. In pollen, pseudouridylated easiRNAs are localized to sperm cells, and we found that *PAUSED/HEN5 (PSD)*, the plant homolog of Exportin-t, interacts genetically with Ψ and is required for transport of easiRNAs into sperm cells from the vegetative nucleus. We further show that Exportin-t is required for the triploid block: chromosome dosage-dependent seed lethality that is epigenetically inherited from pollen. Thus, Ψ has a conserved role in marking inherited small RNAs in the germline.

**One-Sentence Summary:** Pseudouridine marks germline small RNAs in plants and mammals, impacting epigenetic inheritance via nuclear transport.

---

The germlines of both plants and metazoans have abundant small RNA, many of which are cell-type specific. In the mammalian germline, spermatids accumulate 26-28nt PIWI-interacting RNAs (piRNAs) which silence TEs, while spermatocytes accumulate 29-30nt piRNA that also target genes^5^. In plants, the male germline terminates in mature pollen, which contains two cell types, the pollen grain (or vegetative cell), and two sperm cells enclosed within its cytoplasm. Plants do not have piRNAs, but instead 21-22nt epigenetically activated siRNAs (easiRNAs) are produced from TEs in the vegetative nucleus (VN) and translocated to the sperm cells^6–8^. Furthermore, 24nt siRNAs from some DNA transposons are translocated into developing meiocytes in the anther from the surrounding layer of somatic cells (the tapetum)^7^. Both types of small RNA are produced by the plant-specific RNA polymerase, Pol IV^7,9,10^ and mediate epigenetic inheritance after fertilization ^9,11^. In maize, 21nt and 24nt “phasiRNAs” are also transported from epidermal and tapetal cells into the meiocytes, demonstrating that these mechanisms may be conserved^12^. Both miRNAs and siRNAs in plants are modified by 2′-O-methylation, while in animals only piRNAs have this modification^13^.

We set out to determine whether pseudouridine (Ψ) was present in small RNAs (Fig. 1). Ψ is the most common non-canonical base in RNA but has not been explored in small RNAs due to difficulties in detection. High-throughput techniques to detect pseudouridylation in longer RNAs specifically label Ψ with N-cyclohexyl-N’-(2-morpholinoethyl)-carbodiimide-metho-*p*-toluenesulfonate (CMC) to block reverse transcriptase (RT) during RNA-seq library preparation^14–17^. However, in the case of small RNA, RT blocks would result in truncated sequences too short to map back to the genome. More recently, anti-Ψ antibody binding^18^, as well as bisulphite treatment at neutral pH^19^ achieve similar results. In mouse, long primary miRNA precursor transcripts (pri-miRNAs) have been shown to contain Ψ^20,21^, so we initially tested for the presence of Ψ in small RNAs from mouse NIH/3T3 cells. Following immunoprecipitation with a Ψ-specific antibody, we could robustly detect Ψ using a combination of mass spectrometry (Extended Data Fig. 1a) and dotblots (Extended Data Fig. 1b) using *in vitro* synthesized pseudouridylated transcripts as a positive control. Using this antibody, we then performed pseudouridine immunoprecipitation (Ψ-IP) and sequencing of small RNAs from NIH/3T3 cells before and after knockdown of the Ψ synthase PUS1 (Extended Data Fig. 1c-e; Supplementary Dataset S1). We detected Ψ in several miRNAs (Extended Data Fig. 1c) including members of the let-7 family which were validated by qPCR (Extended Data Fig. 1d). let-7 was unaffected by *PUS1* knockdown, however, consistent with its interaction with DKC1 and TruB1 instead^20,21^. But four other pseudouridylated miRNA were strongly impacted in the *PUS1* knockdown (Extended Data Fig. 1e; Supplementary Dataset S1).

**Fig. 1.**
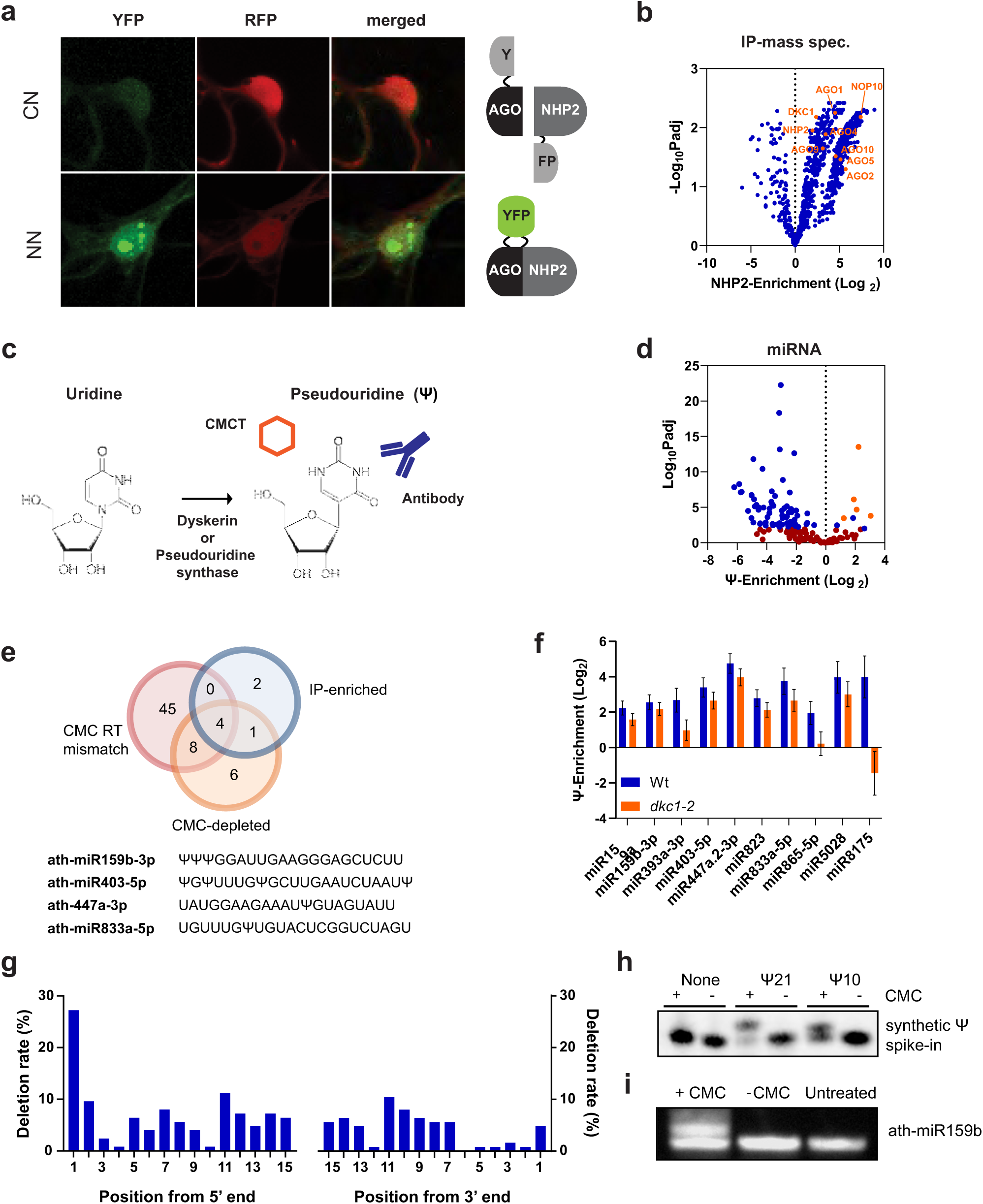
microRNAs are pseudouridylated in plants and mammals. (**a**) Bimolecular fluorescence complementation confirming interaction of AGO3 with the Dyskerin co-factor NHP2 in Arabidopsis (Methods). Split YFP was fused to the C- and N-, or N- and N-terminals of AGO3 and NHP2. Reconstituted YFP signifies interaction. p35S::RFP acts as a transformation control. (**b**) Volcano plot showing proteins co-purified with GFP-NHP2 as compared to IPs performed in wild-type plants. H/ACA ribonucleoprotein complex subunits and AGO family proteins are depicted in orange (p-value adjusted by Benjamini-Hochberg correction). (**c**) Structure of uridine and pseudouridine (Ψ), enzymes involved in catalysis, and methods of detection with specific antibodies and chemical modification with CMC (Methods; Supplemental Fig. 2). (**d**) Volcano plot of miRNAs enriched by Ψ-IP compared to unbound fractions from *Arabidopsis* flower buds (DESeq2). Blue, p<0.01; orange, both Ψ-IP-enriched and depleted from libraries following CMC-treatment (p<0.01); dark red, not significant. (**e**) Venn diagram showing overlap of miRNAs detected in flower buds by each technique. Predicted Ψ sites in miRNAs detected by all 3 techniques. (**f**) miRNAs enriched by Ψ-IP in WT and *dkc1* mutants in Arabidopsis flower buds (**g**) Frequency of miRNA modification at each site based on its proximity to the 5′ or 3′ end based on CMC/Mn^2+^ sequencing (**h**) Northern blots of three synthetic 21mers (sequence UGACACAGGACUACGGACGUAU) either unpseudouridylated, or pseudouridylated at position 10 or 21, treated (+) or mock treated (-) with CMC, and probed with a matching DIG-labeled probe. (**i**) Flower bud small RNA treated (+), mock treated (-) with CMC, or untreated (no mock treatment) and probed with miR159b to reveal mobility shifts.

In search of factors that might modify small RNAs in plants, we performed a yeast two-hybrid screen for interactors of *Arabidopsis* ARGONAUTE 3 (AGO3) using a cDNA library of transcripts from *Arabidopsis* flowers^22^. The screen (Methods) identified a potential interactor of AGO3, namely H/ACA RIBONUCLEOPROTEIN COMPLEX SUBUNIT 2 (NHP2), and we confirmed the interaction by bimolecular fluorescence complementation in phase-separated nucleolar bodies (Fig. 1a). NHP2 plays a key role in RNA pseudouridylation via the pseudouridine synthase Dyskerin (DKC1), often, though not always, via snoRNA guides^23–25^. NHP2 was also in close proximity with other AGO proteins, notably AGO4, which has been found in nucleolar bodies and in related Cajal bodies (Extended Data Fig. 1f)^26^. Immunoprecipitation of GFP-tagged NHP2 followed by mass spectrometry revealed interactions with other subunits of the Dyskerin complex (DKC1 and NOP10) as well as components of the RNAi machinery, including AGO1, 2, 4, 5, 9 and 10 (Fig. 1b; Supplementary Dataset S2). Immunoprecipitation of GFP-tagged NHP2 followed by RNA sequencing revealed snoRNAs, snRNAs and several pri-miRNAs that co-immunoprecipitated with NHP2 either directly or as part of a larger complex (Supplementary Dataset S3; Extended Data Fig. 1g). In order to determine whether pri-miRNAs were pseudouridylated in plants, as they are in animals, we then performed Ψ-IP and sequencing on mRNAs from *Arabidopsis* flower buds (collectively called the inflorescence). Several pri-miRNA transcripts were enriched, and there was a tendency for those enriched by immunoprecipitation with GFP-tagged NHP2 to also be enriched by Ψ-IP (p<0.01, ANOVA) (Extended Data Fig. 1h; Supplementary Dataset S4).

Encouraged by these results, we tested three approaches to detect Ψ directly in small RNAs in *Arabidopsis* (Methods; Fig. 1c; Extended Data Fig. 2a): (i) Ψ-IP followed by small RNA sequencing, (ii) CMC treatment of small RNA libraries following 5ʹ-adapter ligation, to block RT and deplete small RNAs that contain Ψ, and (iii) reverse transcription of CMC-treated small RNAs in the presence of Mn^2+^, which causes deletions and mutations at Ψ residues instead of a reverse transcription stop^27^. To determine the efficacy of these techniques, we examined coverage at known pseudouridylation sites in rRNA and tRNA. Ψ at UNΨAR motifs is directed by PUS7 and was detected by CMC-depletion in WT but not in *pus7* mutants (Extended Data Fig. 2b, c). Ψ at position 55 in tRNA depends on PUS10 (Extended Data Fig 2b). tRNA fragments containing Ψ- 55 were IP-enriched and CMC-depleted, contained mismatches in CMC/Mn^2+^-treated libraries, and were ablated in *pus10* mutants (Extended Data Fig 2b, d, e). Predicted sites of rRNA pseudouridylation were also enriched by Ψ-IP, depleted by CMC treatment, and mismatched in CMC/Mn^2+^ libraries (Extended Data Fig 2f-i). Thus, all three methods robustly detected Ψ in small RNA fragments derived from tRNA and rRNA.

We then applied our assays to detect Ψ in mature small RNAs from *Arabidopsis* inflorescence and detected more than fifty pseudouridylated mature miRNAs (Fig. 1g), of which four (miR159b-3p, miR403-5p, miR447a-3p and miR833a-5p) were detected using all three assays (Fig. 1d,e; Supplementary Datasets S5-7). *pus7* and *pus10* mutants had no detectable effect on Ψ-IP enrichment or CMC-depletion of miRNA, but we expected that DYSKERIN (DKC1) might be responsible, given the enrichment for pri-miRNA in pulldowns with the Dyskerin co-factor NHP2 (Extended Data Fig 1b). As mutants of NHP2 and other members of the DKC complex are lethal^28^, we rescued *dkc1* knockout mutations with an embryo-specific DKC1 cassette (Extended Data Fig. 3a), which restored about 10% of wild-type *DKC1* expression in the leaves (Extended Data Fig. 3b) and allowed seeds to germinate into viable but infertile plants (Extended Data Fig. 3c). Several miRNAs were significantly enriched by Ψ-IP in WT leaves (Fig. 1f) of which 3/10 required DKC1 for pseudouridylation (Fig. 1i). We conclude that just as some miRNAs are pseudouridylated by PUS1 in the mouse (Extended Data Fig. 1h), some are pseudouridylated by DKC1 in Arabidopsis, but in both cases multiple pseudouridine synthases must be responsible for miRNA pseudouridylation, just as they are for mRNA pseudouridylation in plants, yeast and mammals^29^.

Ψ-detection by mismatch sequencing revealed that pseudouridylation in *Arabidopsis* miRNAs (Fig. 1g) was prevalent at the 5’ nucleotide, which is frequently uridine in miRNAs and contributes to AGO loading preference^30–32^. Similarly, siRNAs detected by Ψ-IP enrichment and CMC-depletion, were enriched for 5′ and 3′ U (Extended Data Fig. 4a,b). As pri-miRNAs associate with NHP2 and contain Ψ (Extended Data Fig. 1g), we predicted that Ψ in mature miRNAs was deposited prior to processing of pri-miRNAs into small RNAs by the endonuclease DICER. cDNA reverse transcribed from pri-miR159b in the presence of CMC and Mn^2+^ revealed extensive mismatches (pseudouridylation) at the 5’ dicing site (25%Ψ), which was exacerbated in a *hyponastic leaves1 (hyl1)* mutant (75%Ψ) that accumulates pri-miRNAs due to loss of the microprocessor component R2D2 (Extended Data Fig. 4c). Ψ was also detected at 5’ and 3’ sites in mature miR159b, suggesting that pseudouridylated pri-miRNAs are processed into mature miRNAs (Extended Data Fig. 4c). We confirmed the presence of Ψ in mature miR159b on Northern blots, where CMC-treatment induces a mobility shift in pseudouridylated small RNAs (Fig. 1h,i). Intriguingly, miR159b is inherited from sperm cells by the fertilized endosperm, where it impacts seed development^3^. As miR159b is heavily pseudouridylated (Fig. 1i), this suggests that pseudouridylated small RNA may be inherited from pollen.

We therefore applied Ψ-IP, CMC and CMC/Mn^2+^ treatments to small RNA isolated from pollen. We found that known sites of pseudouridylation in 18S rRNA were robustly detected in rRNA fragments in pollen by all three techniques (Extended Data Fig. 5a-d). In both CMC and Ψ-IP datasets, miRNAs in pollen had a much greater level of pseudouridylation than in flower buds (Fig. 2a; Supplementary Datasets S7-9). This effect was even more dramatic for siRNAs, which were heavily pseudouridylated in pollen (Fig. 2b). Pseudouridylated siRNA (Ψ-siRNA) were associated with most annotated TE families, although there were notable exceptions (e.g. AtGP2N and AtGP2). Next, we tested whether Ψ-miRNA and Ψ-siRNA were enriched in sperm cells purified by FACS, or were immunoprecipitated with AGO proteins from total pollen (Fig. 2c). AGO 2, 5, 6, 7 and 9 bind siRNA, and are restricted to sperm cells in mature pollen^33^, where AGO5 and AGO9 are localized to the perinuclear space (Fig. 1d,e). AGO1 on the other hand binds mostly miRNA and accumulates in both SC and VN^33^.

**Fig. 2.**
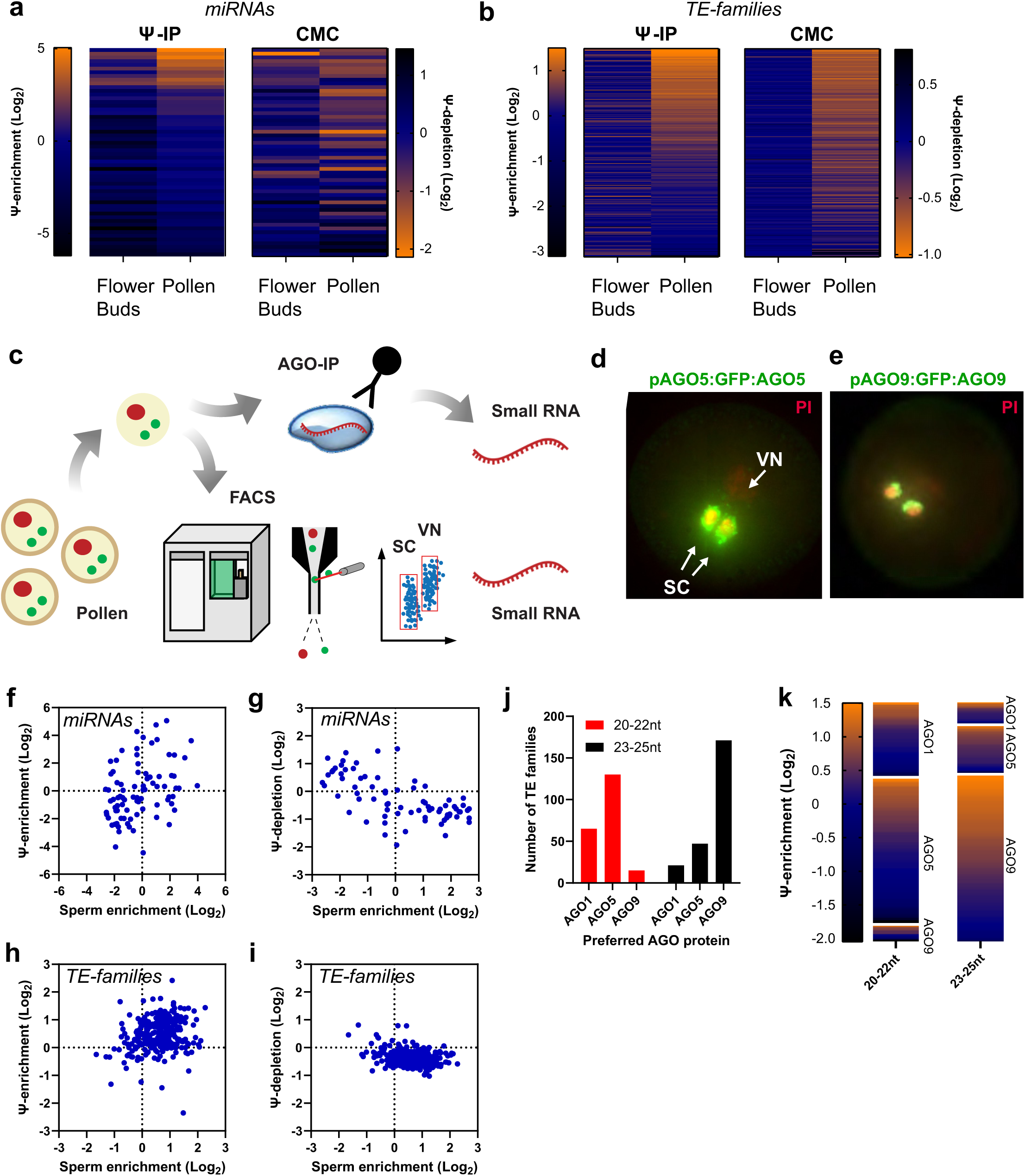
Pollen small interfering RNAs are highly pseudouridylated and loaded into ARGONAUTE proteins in sperm cells. (**a**) Heatmaps of individual miRNA Ψ-enrichment in flower buds and pollen by Ψ-IP (Log_2_ IP/Unbound; n = 5) and by CMC-mediated depletion from small RNA libraries (Log_2_ CMC+/CMC-; n = 2) assessed by DESeq2. (**b**) Heatmaps of siRNA Ψ-enrichment from individual TE families in flower buds and pollen by Ψ-IP and CMC-depletion as in (a). Small RNAs from most TE families were enriched for Ψ in pollen. (**c)** Small RNAs were sequenced from sperm cells isolated from pollen grains by FACS, and by RNA immunoprecipitation (RIP-seq) with antibodies to ARGONAUTE (AGO) proteins. (**d**) GFP::AGO5 fusion protein expressed from *AGO5* promoter (*pAGO5*) in mature pollen is found in the perinuclear space around sperm cells (SC). DAPI was used to stain the vegetative nucleus (VN). Reproduced with permission from Borges *et al.*^34^ (**e**) AGO9::GFP fusion protein expressed from the *AGO9* promoter (*pAGO9*) in mature pollen. (**f-i**) Association of pseudouridylation determined by Ψ-IP (f, h) and CMC-depletion (g, i) in pollen with sperm-cell abundance by FACS for miRNAs (f,g) and siRNAs from TE families (h,i)^6,34^. (**j**) Bar chart showing number of preferred TE-families for each AGO protein split by siRNA size (20-22nt; 23-25nt). Each TE-family was assessed for enrichment in AGO1, AGO5 and AGO9 IP/input for two size classes (20-22; 23-25nt), with the highest enrichment signaling a preference for a certain AGO. (**k**) Heatmaps showing siRNA Ψ-enrichment from individual TE families split by AGO-preference and size-class.

To determine whether pseudouridylation of small RNA reflects their localization, we compared pseudouridylation of pollen miRNAs to sperm cell localization. As expected from previous studies^34^ and consistent with AGO1 localization, miRNA were found in both SC and pollen, and some miRNA in each case were pseudouridylated (Fig. 2f,g). Consistent with the localization of AGO 2, 5, 6, 7, and 9, however, siRNA were strongly enriched in sperm cells, and siRNA from most TE-families were pseudouridylated (Fig. 2h,i). To determine whether individual AGO proteins preferentially bound pseudouridylated siRNA, we performed small RNA sequencing after immunoprecipitation (RIP-seq) with anti-AGO9 antibodies (Methods) and compared the abundance of siRNA of each size class in each TE family with published RIP-seq data from AGO1 and AGO5^35^, with AGO5 and AGO9 being preferred for the 20-22nt and 23-25nt size classes, respectively (Fig. 2j). Next, we examined pseudouridylation levels of these families by sequencing following Ψ-IP to determine if AGO-binding influenced pseudouridine levels, and we found that all AGO proteins bound pseudouridylated siRNA from a similar proportion of TE families (Fig. 2k). This was consistent with association of the pseudouridylation machinery with all of these argonautes by IP-MS (Fig 1b). We therefore sought to determine what other factors might control pseudouridylation in pollen.

Pseudouridylation has been implicated in cytoplasmic localization of mRNA and tRNA suggesting a role for Ψ in small RNA transport^16,36^. Pol IV-dependent 21-22nt easiRNAs in pollen are also mobile and move into sperm cells from the VN^6–8^. To further investigate a role for Ψ in inter-cellular mobility of small RNAs, we probed the function of Ψ in sperm cell localization. Since our *dkc1* mutant was completely sterile (Extended Data Fig. 3c), we could not use this to investigate the function of Ψ in pollen. Additionally, other PUS enzymes likely have considerable redundancy, as twenty exist in Arabidopsis^29^. Pseudouridylation of tRNAs by Pus1^+^ contributes to nuclear export in yeast, and genetically interacts with Loss of Suppression 1 (Los1^+^), which encodes the tRNA export factor Exportin-t^36^. Exportin-t binds the upper minihelix of L-shaped tRNA, consisting of the acceptor-stem and the TψC-loop, and terminating at Ψ55 and C56^37^. This minihelix includes the 22nt 3′ tRNA fragment with a 5′ Ψ and 3′CCA, resembling a 21-22nt small RNA duplex^38^. Intriguingly, the *Arabidopsis* homolog of *Los1*^+^, which is known as *PAUSED/HEN5 (PSD)*, was previously identified in genetic screens for mobile small RNA phenotypes, along with mutants in *ARGONAUTE7* (*zippy*) and *HASTY (hst*). *HST* encodes Exportin-5 and has recently been shown to be required for long-range transport of artificial miRNA in Arabidopsis^39–42^. We therefore investigated whether *psd* mutants could be used as a proxy for the function of Ψ in siRNA, as mutants are viable and fertile.

First, we demonstrated that the *psd* mutation in *Arabidopsis* is synthetically lethal with *pus7* in a manner similar to *los1*^-^ and *pus1*^-^ from yeast. We found that *pus7-/-*; *psd+/-* double mutants were semi-sterile (Extended Data Fig. 6a). In contrast, *pus7+/-; psd-/-* plants had fertile pollen but had 25% aborted seed compared with less than 5% in *psd* alone (Extended Data Fig. 6b,c). Viable double homozygous seeds were not recovered from either genotype upon self-fertilization. These results indicated that tRNA require *PUS7* for modification in diploid cells before meiosis (*pus7-/-*), and subsequently require *PSD* for nuclear export in haploid pollen (from *psd+/-* parents). Next, we performed Ψ-IP and CMC depletion on small RNA from WT and *psd* mutant pollen as well as WT and mutant inflorescence (flower buds) and found robust detection of pseudouridylation sites in rRNA (Extended Data Fig. 5a-f). We sought to determine if *psd* had an effect on pseudouridylation of miRNA and siRNA in inflorescence and pollen. Of those miRNAs that were significantly pseudouridylated in the inflorescence, only miR403 lost Ψ in the mutant (Extended Data Fig. 5g), while miRNA overall were unaffected (Extended Data Fig. 5h). Similarly, siRNA from TEs were mostly unmodified in the inflorescence, and were unaffected by *psd* (Extended Data Fig. 5i). In pollen, however, miRNA were modestly depleted of Ψ in *psd* relative to WT (Fig. 3a; p<0.01 ANOVA) while easiRNA were strongly depleted of Ψ (Fig. 3b). easiRNA biogenesis requires both Pol IV and the most abundant miRNAs in pollen, miR845a and miR845b, which target LTR retrotransposons at their reverse transcription primer binding site^9,35^. 20-22nt easiRNAs that depend on miR845b were heavily pseudouridylated, and this depended on *psd*, as did pseudouridylation of 20-22 nt easiRNAs and 23-25nt siRNAs that depend on Pol IV (Fig. 3b; Extended Data Fig. 7a). However, Pol IV-independent siRNA had very little Ψ which was unaffected in *psd* (Fig. 3b; Extended Data Fig. 7a).

**Fig. 3.**
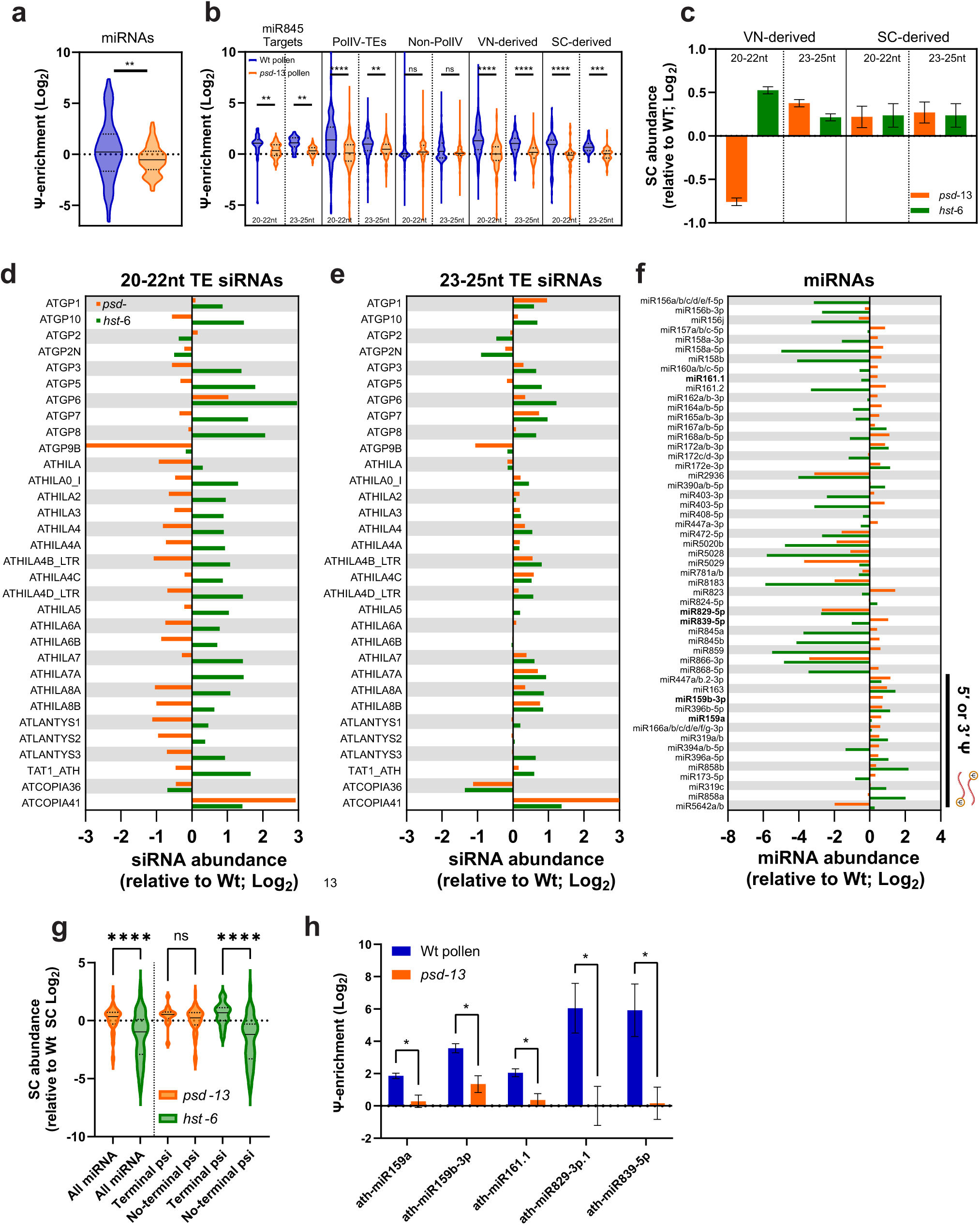
Transport of pseudouridylated siRNAs into sperm cells depends on *PAUSED* (Exportin-t). (**a**) Ψ-enrichment of miRNAs in wild-type and *psd*-13 pollen. A modest reduction in Ψ-enrichment of miRNAs was observed in *psd* pollen (p<0.01, paired t-test; n = 3). (**b**) Ψ-enrichment of siRNAs was determined by Ψ-IP and small RNA sequencing in wild-type and *psd-13* mutant pollen. Individual TEs belonging to each group (miR845 targets, PolIV/Non-PolIV and SC/VN-derived siRNA) and size class (20-22nt or 23-25nt) are plotted based on their enrichment by Ψ-IP. Asterisks indicate significance (p<0.01 (**), 0.001(***), 0.0001 (****), ANOVA with Bonferroni correction for multiple comparisons, n = 3 biological replicates). (**c**) siRNA abundance was determined by sequencing small RNA from FACS sorted wild-type, *psd*-13 and *hst*-6 sperm cells. VN-derived 20-22nt siRNA were depleted from *psd* sperm cells but enriched in *hst* sperm cells relative to wild-type sperm, while 23-25nt, and SC-derived siRNA were at levels similar to wild-type. (**d,e**) Abundance of highly prevalent TE families (Gypsy and two Copia elements) in *psd* sperm and pollen and *hst* sperm relative to wild-type for 20-22nt (d) and 23-25nt (e). (**f**) Abundance of highly prevalent miRNAs (RPM>100) in *psd* sperm and pollen and *hst* sperm relative to wild-type. miRNA with Terminal pseudouridine are indicated with a black bar; miRNA presented in (h) are shown in bold. (**g**) Abundance of miRNA with and without terminal Ψ in *psd* and *hst* mutants relative to wildtype in sperm cells sorted by FACS. miRNA without terminal Ψ were lost in *hst*, but not in *psd* sperm cells (p<0.001; ANOVA, n=3). (**h**) Histogram of miRNA Ψ-enrichment in wild-type and *psd-13* pollen for those miRNA dependent on *PSD* for pseudouridylation (p<0.05; student’s t-test n = 3).

In pollen, Pol IV is prominently expressed in the VN^43^, suggesting that this is where pseudouridylation likely takes place. However, easiRNAs accumulate in sperm cells, and are thought to be transported there from the VN^6–8^. In a recent study, small RNA biogenesis in the VN and in the SC were independently suppressed by expressing the RNAseIII-like protein RTL1 in each cell type^44^. We found that both VN-derived and SC-derived small RNA were enriched for Ψ, and that enrichment depended on *psd* (Fig. 3b; Extended Data Fig. 7b). To determine if PSD affected siRNA transport from VN into sperm cells, we isolated sperm cells from mutant *psd-13* (Exportin-t), mutant *hst-6* (Exportin-5), and wild-type pollen by FACS and compared the abundance of siRNAs and miRNAs (Fig. 3c-e; Extended Data Fig. 7c-l). Strikingly, 21-22nt easiRNAs were reduced in *psd* SC and elevated in *hst* SC, but only those that were derived from the VN, and not for those derived from the SC (Fig. 3c,d). In contrast, 23-24nt siRNA abundance in SC was unaffected by the *psd* and *hst* mutants (Fig. 3c,e). However, both PolIV and non-PolIV TE-siRNA were reduced in *psd* sperm cells, suggesting that pseudouridine is not strictly required for transport (Extended Data Fig. 7c)

HST (Exportin-5) is known to be required for the export and accumulation of miRNA^45^, and miRNA levels in SC were significantly reduced in *hst* mutants (Fig. 3f,g). As expected, most miRNA in sperm cells were unaffected by *psd* (Fig. 3f,g), but individual miRNA including miR159 and miR403 significantly lost Ψ (Fig. 3h). A likely explanation is that miRNA that lose Ψ in *psd* mutants can use the *HST* pathway instead to accumulate in sperm cells. One prediction of this model is that Ψ-miRNA might be able to use the PSD pathway in *hst* mutants. This prediction was confirmed, as miRNA without 5′ terminal Ψ were lost in sperm cells from *hst* mutants, while those with terminal Ψ were retained, suggesting PSD was sufficient for their transport (Fig. 3f,g; Supplementary Dataset S7). Thus *PAUSED/EXPORTIN-T* is required for transport of pseudouridylated easiRNA into sperm cells from the VN, while *HASTY/EXPORTIN-5* transports unmodified miRNA. This novel function for *PAUSED/EXPORTIN-T* is pollen-specific, as are most pseudouridylated miRNAs and easiRNAs, which are unaffected by *psd* in the inflorescence (Extended Data Fig. 5g-i). It is noteworthy that tRNA have been found to carry a strong signal for intercellular transport in plants, so that mRNA-tRNA fusions are mobile^46^. It is tempting to speculate that Ψ and *PSD* may be involved in intercellular transport of tRNA fusions.

easiRNAs from pollen target imprinted genes in Arabidopsis^9,11^. Imprinted genes in plants are expressed in the seed from either the maternal or the paternal genome and are known as maternally expressed and paternally expressed genes (MEGs and PEGs), respectively. *DEMETER* (*DME*) encodes a DNA glycosylase that is expressed in the endosperm and in the pollen VN, where it removes DNA methylation from TEs upstream and downstream of imprinted genes resulting in parent-of-origin expression^47^. We therefore examined Ψ in siRNAs from DME targets and imprinted genes. We detected Ψ in 21-22nt easiRNAs and 24nt siRNAs from DME targets (Fig. 4a), and from TEs surrounding maternally- and paternally-expressed imprinted genes (MEGs and PEGs; Extended Data Fig. 7m). Pseudouridylation of siRNAs from upstream and downstream of DME targets was lost in *psd-13* mutant pollen (Fig. 4a), and there was a striking depletion of easiRNA from the promoters of DME target genes in *psd* sperm cells relative to WT (Fig. 4b,c). Ψ loss was more significant in MEGs rather than PEGs (p<0.05; Extended Data Fig. 7m) and also affected sperm cell localization of siRNAs from both MEGs and PEGs, with 20-22nt siRNAs depleted and 23-25nt siRNAs enriched in *psd-13* sperm cells compared to wild-type (Extended Data Fig. 7n).

**Fig. 4.**
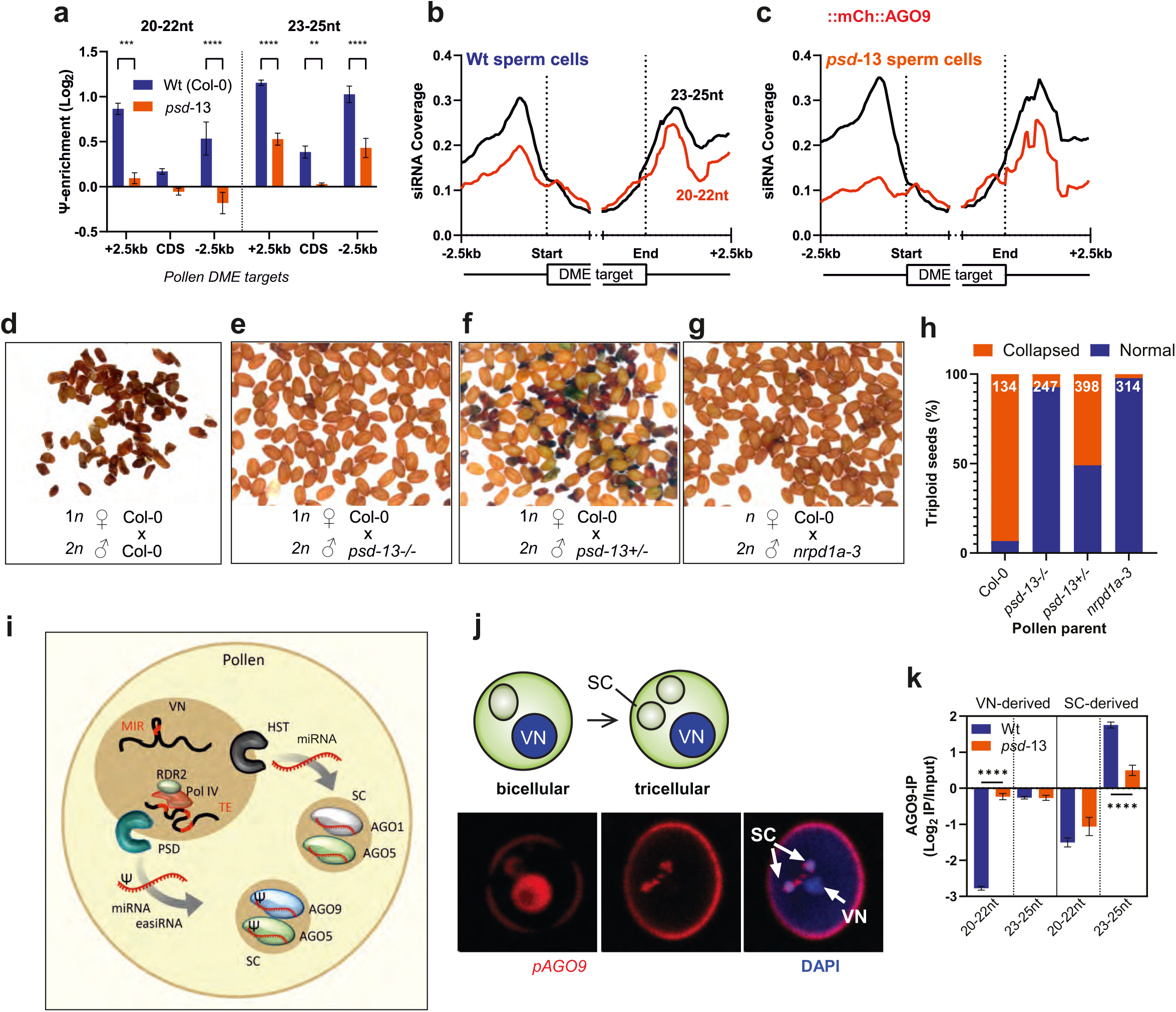
Pseudouridine and Exportin-t mediate epigenetic inheritance and the triploid block. (**a**) Ψ-enrichment (Log_2_ Ψ-IP/unbound) of 20-22 nt siRNAs and 23-25 nt siRNAs surrounding imprinted genes targeted by *DEMETER (DME),* in wild-type (blue) and *psd-13* (orange) pollen (Asterisks indicate student’s t-test; p<0.01 (**), 0.001 (***) and 0.0001(****); n=3). (**b, c**) sperm cell abundance of 20-22 nt siRNAs (red) and 23-25 nt siRNAs (black) surrounding imprinted genes targeted by *DEMETER (DME),* in wild-type (b) and *psd-13* (c) pollen. (**d-g**) Images of representative seeds from crosses between wild-type Col-0 female and (d) *osd1* diploid pollen, (e) double mutant *osd1*; *psd-13-/-* diploid pollen, (f) double mutant *osd1*; *psd-13+/-* diploid pollen (g) double mutant *osd1; nrpd1a-3* diploid pollen. *NRPD1*a encodes the large subunit of PolIV. (**h**) Frequency of aborted seeds from the same crosses. (**i**) Model for transport of small RNA during pollen development. As the pollen matures 21/22 nt easiRNAs and 24nt siRNAs are produced by Pol IV in the vegetative nucleus and pseudouridylated along with some miRNAs. pseudouridylated miRNAs and easiRNAs are exported via PSD (Exportin-t) into sperm cells, where they are loaded onto AGO1, AGO5 and AGO9, while unmodified miRNAs are transported via HST (Exportin-5) and loaded onto AGO5 and AGO1. Pseudouridylated sperm cell easiRNAs mediate dose-dependent lethality (the triploid block) by targeting maternally expressed imprinted genes in the seed. (**j**) mCherry::AGO9 fusion protein was expressed from the *AGO9* promoter (*pAGO9*) in developing pollen. DAPI was used to stain nucleic acids (blue). AGO9 is present in the VN at the binuclear stage but is enriched in the sperm cells in mature pollen. (**k**) AGO9 IP from wild-type and *psd*-13 pollen shows increased binding of VN-derived 20-22nt siRNA and a decrease in 23-25nt SC-derived siRNA to AGO9 in *psd*-13.

Pol IV-dependent 21-22nt easiRNAs and miR845b mediate the triploid block, a dose-dependent epigenetic phenomenon whereby *Arabidopsis* seeds abort during development, if they inherit an excess of paternal genomes from diploid instead of haploid pollen^9–11^. As 21-22nt Pol IV-dependent easiRNAs are lost from sperm cells in *psd-13* pollen, this suggested that *psd-13* might rescue triploid seed abortion^9^. As expected, we observed ∼90% seed abortion when WT Columbia plants were crossed with *omission of second division 1 (osd1)* mutants, which have diploid pollen (Fig. 4d). However, when *osd1*;*psd-13*-/- diploid pollen was applied, this resulted in only ∼10% seed abortion, similar to when *osd1*; *nrpd1a* diploid pollen was applied, which is deficient in the large subunit of Pol IV (NRPD1; Fig. 4e-h). Levels of miR845a and miR845b were not affected in *psd-13* mutant pollen, and were not responsible for rescuing the triploid block (Extended Data Fig. 7o)^9^. Importantly, *osd1; psd/+* pollen parents gave rise to 50% aborted seeds (Fig. 4f,h). One possible explanation is that *psd* acts gametophytically, within the pollen grain itself, supporting the observation that VN-derived 21-22nt easiRNA, but not SC-derived small RNA, are required for the triploid block^44^.

Exportin-t is therefore essential for the triploid block and for pseudouridylation of easiRNAs. This implicates Ψ as a key modification in cell-cell transport of small RNAs in the germline via Exportin-t (Fig. 4i) and for epigenetic inheritance in the form of the triploid block. Cytoplasmic connections between VN and sperm cells are thought to be involved^6,8^, consistent with a role for nucleocytoplasmic exportin. AGO5 is found in these “male germ units”^34^, and AGO5 is also required at least in part for the triploid block^35^. AGO9 has a similar localization (Fig. 2d,e) and has previously been associated with siRNA movement in pollen^48,49^. We therefore examined AGO9 localization using an mCherry N-terminal fusion^50^ and found that AGO9 was first localized in the VN at the bicellular stage, but then moved to the sperm cells by the tricellular stage (Fig. 4j). This localization is consistent with a role for AGO9 before, and after siRNA transport as previously proposed^48,51^. We therefore performed immunoprecipitation of small RNA from WT and *psd* pollen using antibodies to AGO9. Remarkably, AGO9-associated 21-22nt easiRNA were substantially more enriched in *psd* mutants than in WT (Fig. 4k). One interpretation is that easiRNA are trapped in the VN in *psd* mutants, where they are subsequently loaded onto AGO9.

easiRNAs and phasiRNAs in the plant germline have similar functions to piRNAs in the mammalian germline, and so we asked if high levels of Ψ were conserved in piRNAs. We used Ψ- IP as well as CMC depletion to detect Ψ in small RNAs from 8-week *post-partum* mouse testis. 3′-tRNA fragments (tRFs) were enriched by Ψ-IP (Extended Data Fig. 8a) and depleted by CMC (Extended Data Fig. 8b). Importantly many piRNAs from TEs, including from LINE and LTR retroelements, were strongly enriched for Ψ (Extended Data Fig. 8c,d). Up to 3% of piRNAs were differentially enriched for Ψ, though many more were immunoprecipitated and hence potentially pseudouridylated (Extended Data Fig. 8d). No individual piRNA cluster was significantly enriched or depleted, suggesting sequence-rather than locus-specificity (Extended Data Fig. 8e,f). 3′-tRF that were depleted by CMC treatment were 22nt in size as expected for 3′-tRF containing Ψ-55 from mature tRNA (Extended Data Fig. 8g) while piRNA were mostly 30nt in size as expected (Extended Data Fig. 8h). The same small RNA sizes and classes were detected by Ψ-IP (Extended Data Fig. 8i,j). Thus, not only 3′-tRFs, but also piRNAs and miRNAs in mammals are pseudouridylated. Intriguingly, 5′-tRFs are also pseudouridylated, and have been reported to be inherited^52,53^.

Thus, germline small RNA in plants and mammals are heavily pseudouridylated, which at least in plants, depends on Exportin-t. This is true of both VN-derived and SC-derived small RNA in pollen, implicating nucleocytoplasmic shuttling in pseudouridylation, as previously proposed^54^. Intriguingly, *PSD* is also required for transport of VN-derived easiRNA to the SC, and for the triploid block. Therefore, either Ψ or intercellular transport or both, are required for epigenetic inheritance in the form of the triploid block. Why are germline small RNAs so heavily modified in both plants and mammals? An intriguing possibility is that modifications of RNA in the germline may avoid viral surveillance systems after fertilization, which could otherwise recognize inherited small RNA as “non-self”^55^. RTL1 is such a viral surveillance mechanism and is induced strongly on viral infection^56^. While VN-derived easiRNA are suppressed when RTL1 is expressed in the VN, they are not suppressed when RTL1 is expressed in SC even though VN-derived easiRNA accumulate in SC^44^. An intriguing explanation might be that easiRNA pseudouridylation requires export from the VN, and RTL1 cannot recognize these easiRNA once they are pseudouridylated.

## Materials and methods

### Cell lines and tissues

NIH/3T3 (ATCC, CRL-1658) were cultured in DMEM (Invitrogen), supplemented with 10% fetal bovine serum (FBS) and 1% penicillin/streptomycin/glutamine. Cells were obtained from ATCC and tested negative for mycoplasma contamination. Testes were harvested from C57Bl/6 mice at 8 weeks *post-partum* and flash frozen in liquid nitrogen.

### Plant growth conditions

*Arabidopsis* plants were grown under long-day conditions at 22°C. Seeds were always surface-sterilized with sodium hypochlorite, sown on Murashige and Skoog medium and stratified for 3 days at 4°C. Seedlings were transplanted to soil 2 weeks after germination and grown under long-day conditions at 22°C. The following mutants were used: *pus7*-1 (SALK_098773), *pus10*¬-1 (SALK_091360), *nrpd1a*-3 (SALK_128428). Homozygous mutants were identified using PCR with primers specific to the predicted site of T-DNA integration (Supplementary Table S1). *psd*- 13 and *hst*-6 alleles were previously identified^39,57^. The *osd1*-3 mutant was kindly provided by Raphael Mercier.

### Yeast two hybrid screening

The two-hybrid screen was performed by Hybrigenics (Paris, France) using an Arabidopsis thaliana Col-0 library (ATFO) prepared from young flowers, using an in-house cDNA clone of AGO3 as bait. 7 candidate proteins were tested by bimolecular fluorescence complementation, leading to the identification of AT5G08180, which encodes the H/ACA ribonucleoprotein complex subunit 2 (NHP2), as a high confidence interactor. This was subsequently confirmed by Mass Spectrometry.

### Bi-molecular fluorescence complementation

BiFC was performed using pBiFC-2in1 vectors^58^. AGO3 and NHP2 were fused in all orientations (NN, CN, NC, CC; only NN and CC complemented), while other AGO proteins were fused only in NN configuration. Tobacco leaves were infiltrated essentially as previously described^59^. Briefly, agrobacterium cultures harboring pBiFC vectors were grown overnight before subculturing in 50 mL of LB broth with antibiotics and 20 µM acetosyringone. Cultures were pelleted by centrifugation, and resuspended in infiltration buffer (10 mM MES, 10 mM MgCl2, 100 µM acetosyringone). Leaves of ∼3 weeks old *Nicotiana benthamiana* plants were infiltrated on the abaxial surface using a syringe without a needle. Three days post-infiltration leaf discs were removed and imaged using a Zeiss LSM780 confocal microscope. Primers for cloning AGO proteins and NHP2 are detailed in Supplementary Table S1.

### Construction of viable *DYSKERIN* (*dkc1*) mutants in Arabidopsis

Dyskerin (*dkc1*) viable mutants were generated as follows. *AtNAP57 (DKC1)* genomic sequence was amplified and cloned into the pENTR vector. The *ABI3* promoter sequence was amplified and inserted using the NotI restriction site. *pABI3:AtNAP57* cassette was transferred to the pGWB501 vector using the Gateway cloning system and used for Agrobacterium-mediated, floral dip Arabidopsis transformation of *nap57*/+ (SALK_031065) heterozygous plants. Homozygous transformant lines were obtained by three rounds of Hygromycin resistance selection followed by verification of *nap57/nap57* genetic background by PCR.

### Co-immunoprecipitation and mass-spectrometry analyses

For each co-IP, 0.6 g of tissue from GFP-NHP2 plants was used. The powder was extracted for 30 min in 1.5 ml of lysis buffer (50 mM Tris HCl pH 8.0, 50 mM NaCl, 1% Triton X-100 and 1× cOmplete™ ethylenediaminetetraacetic acid (EDTA)-free protease inhibitor (Roche). After removal of cell debris by centrifugation (twice 10 min, 16 000 g, 4^◦^C) the cleared supernatants were incubated for 30 min with anti-GFP antibodies coupled to magnetic microbeads (µMACS GFP isolation Kits, Miltenyi). Beads were loaded on magnetized MACS separation columns equilibrated with lysis buffer and washed four times with 300 µl washing buffer (50 mM Tris HCl pH 7.5, 0.1% Triton X-100). Samples were eluted in 100 µl of pre-warmed elution buffer (Miltenyi). Control IPs were performed using Col-0 plants and GFP antibodies. Eluted proteins were digested with sequencing-grade trypsin (Promega). Peptide mixtures were separated using Evosep One (Evosep Biosystems) chromatography coupled to Orbitrap Exploris 480 (Thermo) at IBB PAS, Warsaw. Data were searched against the TAIR10 database. Peptides were identified with the Mascot algorithm (Matrix Science, London, UK). The total number of MS/MS fragmentation spectra was used to quantify each protein from three replicates. The statistical analysis based on spectral counts was performed using a homemade R package that calculates fold change and p values using the quasi-likelihood negative binomial generalized log-linear model implemented in the edgeR package^60^. P-value was adjusted using the Benjamini–Hochberg method from the stats R package. The size factor used to scale samples was calculated according to the DESeq2 normalization method^61^.

### miRNA precursor pull-down with NHP2-GFP

Anti-GFP antibodies were used to immunoprecipitate NHP2-GFP from three independent transgenic lines expressing GFP-NHP2 under native promoter and WT plants as a negative control. RNA was converted into cDNA libraries using NEBNext Ultra II Directional RNA Library Prep Kit for Illumina. Libraries were sequenced with Nextseq HO 100bp paired end. Adapters and unpaired reads were removed using Trimmomatic and files were analyzed using Salmon software. Obtained TPM values for each transcript were adjusted with 0.1 or 0.01 for snRNA/snoRNA and pri-/pre-miRNA respectively to avoid dividing by zero in the case of non-detected transcripts. IP values were normalized against input, and Fold Change was calculated for GFP-NHP2 vs. negative control (GFP IP in WT plants). Reproducibly enriched RNAs in NHP2 IP had FC ≥ 1.2 in at least two lines.

### RNA extraction of mammalian cells

Cells were lysed in Qiazol (QIAGEN) and total RNA was purified using the miRNEasy mini kit (QIAGEN) according to the manufacturer’s instructions. Tissues were lysed in Qiazol (QIAGEN) and total RNA was purified using Tissuelyser II (QIAGEN), as per the manufacturer’s instructions. Small RNAs (<200nt) size-fractionated using RNA Clean & Concentrator 5 column kits (Zymo), as per the manufacturer’s instructions. Testes of 8 weeks old mice were ground under liquid nitrogen and RNA was extracted using Trizol (Thermo Fisher Scientific) according to the manufacturer’s instructions but using 80% EtOH during the wash step.

### Dot-blot

For dot-blot analysis, input RNA or RNA immunoprecipitated with either anti-Ψ or isotypic non-specific antibodies was spotted onto a nitrocellulose membrane and UV cross-linked at 254 nm (120 mJ/cm2). The membranes were blocked in Denhart’s solution (1% Ficoll, 1% polyvinylpyrrolidone, 1% bovine serum albumin; Thermo Fisher) for 1 h at room temperature and incubated with Ψ antibody for 1 h at room temperature. Signal was detected using HRP conjugated secondary antibodies and ECL (GE Healthcare) and developed on a Chemidoc MP machine (BioRad).

### Pseudouridine Immunoprecipitation

Small RNAs were isolated from 0.01-0.1 mg total RNA, first by separating <200 nt RNAs with the Zymo clean and concentrator kit, and subsequently gel extracting RNA 20-30 nt on a 15% polyacrylamide gel. Small RNAs were eluted from the crushed gel slice overnight in 500 µL RNA elution buffer (0.3M NaCl, 0.5 mM EDTA). RNA was resuspended in 50 µL water, typically containing ∼4-40 ng/µL RNA. Resuspended RNA was mixed with 200 µL IPP buffer (150 mM NaCl, 0.1% NP-40, and 10 mM Tris-HCl [pH 7.4] + 10 U/mL RNAse inhibitor) and 5 µg of mouse monoclonal pseudouridine antibody (D347-3, MBL intl.). Samples were incubated for 2 hours at 4°C with rotation before adding 50 µL sheep anti-mouse IgG dynabeads washed in IPP buffer (Invitrogen). The unbound fraction was retained, and the beads were washed 4 times with IPP buffer. To prepare RNA from the unbound fraction, 400 µL binding buffer from the Zymo clean and concentrator kit was added, and RNA was run through the column according to manufacturer’s instructions. To elute bound plant siRNA from the beads, 150 µL Trizol was added and incubated for 2 minutes before placing tubes on a magnetic rack. Trizol was transferred to a new tube and 30 µL chloroform was added and mixed thoroughly. Samples were centrifuged for 10 minutes at maximum speed and the aqueous phase was pipetted into a new tube with an equal volume of 100% ethanol. Mouse small RNAs were eluted using 6.7 mM Ψ triphosphate in IPP buffer with RNase-OUT (Thermo Fisher Scientific) for 30 minutes at 37°C. Samples were then run through a Zymo Clean and Concentrator kit according to manufacturer’s instructions and eluted in 8 µL. Libraries were prepared using NEBNext small RNA sequencing kit (New England Biolabs) or NextFlex small RNA sequencing kit (Perkin Elmer) according to manufacturer’s instructions, with the same temperature modifications as the CMC-treated samples. Libraries were pooled and run on NextSeq 500, HiSeq 4000 or MiSeq high-throughput sequencing systems (Illumina).

### Nucleoside analysis of RNA

RNA samples were incubated overnight with a 2x nuclease mix containing 62.5 units of Benzonase (Sigma Aldrich), 5 units of Antarctic Phosphatase (NEB) and 10 mU/µL of phosphodiesterase I (PDEI) from Crotalus adamanteus venom (Sigma Aldrich) made up in a 5X digest buffer composed of 20 mM Tris-HCl (pH 8), 20 mM MgCl2 and 100 mM NaCl. Samples were then further purified to remove protein by filtration at 14,000 rpm for 30 minutes through Amicon® Ultra-0.5 spin columns (Sigma Aldrich) which have a 30 kDa molecular weight cutoff^62^. Samples were then analyzed using a Q Exactive HF Orbitrap mass spectrometer (Thermo Scientific). Briefly, 2 ul of digested sample was loaded onto a 100Å, 1.8 µm, 2.1 mm X 100 mm ACQUITY UPLC HSS T3 Column (Waters) in an aqueous buffer consisting of 0.1% MS-grade formic acid in water: acetonitrile 98:2 at a constant flow of 0.3ml/min. A gradient of organic solvent was used to facilitate separation and elution as follows: from 2 to 8 minutes the percentage of organic solvent (0.1% formic acid in acetonitrile) increased from 2% to 10%. From 8 to 13 minutes the gradient increased from 10% to 98% organic solvent, where it was held for 1 minute before returning to 2% organic to re-equilibrate the column for the next sample. Samples were read on a full scan positive mode from 100-600 m/z and data were analyzed using Xcalibur software (Thermo Scientific). All nucleosides were compared to exogenous standards which allowed the interpolation of concentrations of sample nucleosides. Isobaric nucleosides were differentiated on the basis of their retention times as compared with pure commercial standards. As an example, Ψ has a retention time of 1.55 minutes in this protocol, whereas uridine elutes at 3.05 minutes.

### RT-qPCR

Cells were lysed in Qiazol (QIAGEN) and total RNA was purified using the miRNEasy mini kit (QIAGEN) according to the manufacturer’s instructions. For mRNA detection, 1 μg of purified total RNA was reverse transcribed using the high-capacity cDNA reverse transcription kit (Applied Biosystems). For specific miRNA quantification, we size-fractionated small RNAs (< 200nt, containing pre-miRNAs) using RNA Clean & Concentrator 5 column kits (Zymo), as per the manufacturer’s instructions. miRNAs were reverse transcribed with miScript II RT kit (QIAGEN). Primers were designed to anneal on the 5p or 3p arm of the miRNA of interest (Supplementary Table S1). miRNAs were quantified using Fast SybrGreen PCR mastermix (Applied Biosystems) according to the manufacturer’s instructions.

### CMC-treatment and small RNA sequencing

For each pair of libraries 4 µg of Arabidopsis inflorescence RNA, prepared using Zymo Quickzol RNA kit, was ligated to the 3′ adapter supplied in the NEB NextSeq kit in a 40 µL reaction. After ligation samples were divided in two and subjected to CMC or mock treatment. 80 µL of 0.5M CMC in BEU buffer (7 M urea, 4 mM EDTA, 50 mM bicine pH 8) was added to 20 µL sample and incubated at 37°C for 30 minutes. Mock treated samples were incubated with 80 µL of BEU buffer. After CMC/mock treatment 25 µL 3M sodium acetate, 3 µL glycogen and 1 mL 100% ethanol was added and samples were incubated for 1 hour at -80C. Samples were centrifuged at 4°C for 30 minutes and the resulting pellets were washed with 80% ethanol, followed by another 1 hour at -80C. Samples were centrifuged at 4C for 30 minutes and the ethanol removed; pellets were allowed to dry for 5-10 minutes before addition of 50 µL freshly prepared 0.05 M carbonate buffer (pH 10.4) containing 4 mM EDTA. Samples were incubated at 37°C for 3 hours and subsequently cleaned using Zymo RNA clean and concentrator kit, eluting in a final volume of 25.5 µL. After cleaning, libraries were prepared according to instructions in the NEBNext small RNA sequencing kit (New England Biolabs) or Nextflex Small RNA-Seq Kit v3 (Perkin Elmer), beginning with annealing of the RT primer, with some modifications. Primer annealing was performed at 65°C for 5 minutes, followed by 37°C for 15 minutes and 25°C for 15 minutes. cDNA synthesis was carried out at 42°C. After PCR amplification libraries were cleaned using Qiagen PCR cleanup kit and run on a 6% acrylamide gel. Bands of ∼140 nt were cut from the gel, crushed, and eluted for 2 hours in DNA elution buffer followed by ethanol precipitation as outlined in the manufacturer’s instructions. Libraries were pooled and run on NextSeq or MiSeq high-throughput sequencing systems (Illumina) at the Cold Spring Harbor Laboratory sequencing facility.

### CMC/Mn2+ treatment and sequencing

For high-throughput siRNA mismatch analysis 2 µg total Arabidopsis RNA was treated as described above for CMC-siRNA sequencing libraries with some modifications. Nextflex Small RNA-Seq Kit v3 (Perkin Elmer) was used for all samples. The RT step was modified to utilize SuperScript II (Thermo Fisher) and a custom 5X First-Strand Buffer containing MnCl2 (250 mM Tris-HCl, pH 8.3; 375 mM KCl; 30 mM MnCl_2_). For pri-miRNA mismatch analysis, 10 µg of total RNA was treated in the same way as for siRNA mismatch analysis. Obtained cDNA was used for amplification with pri-miRNA specific primers. PCR products were gel-purified and cloned into pGEM-T Easy (Promega) for Sanger sequencing. The same number of clones was sequenced for corresponding CMC+ and CMC-samples.

### Northern Blotting

50 µg total RNA was treated with 0.5 M CMC in BEU buffer (typically 25 µL RNA in 200 uL buffer). Samples were cleaned and prepared in the same way as for small RNA sequencing, except that RNA <200 nt was prepared using Zymo RNA Clean and Concentrator to enrich small RNA in a very low volume. Samples were briefly denatured at 65°C for two minutes (higher temperatures should be avoided as these can remove the CMC adducts) before loading on a 15% denaturing polyacrylamide TBE gel. CMC+/CMC-were treated identically, except for addition of CMC. Untreated RNA was loaded directly on the gel after denaturation. Gels were run until the bromophenol blue band reached the bottom. RNA was transferred to Hybond NX membrane using a semi-dry electroblotter for 30 minutes. Crosslinking was performed with EDC^63^. Membranes were blocked using DIG easy-hyb buffer (Roche) followed by incubation with 10 pmol/mL 5′/3′ DIG labeled LNA probe (Exiqon) in easy-hyb buffer at 65°C overnight. Probes were detected using Roche DIG wash and block buffer set, anti-DIG antibody and CDP star reagent according to manufacturer’s instructions.

### Immunoprecipitation of AGO9

Immunoprecipitation of AGO9 from Arabidopsis pollen was performed as previously described^64^. Pollen was collected from open Arabidopsis flowers as described above, prior to grinding in liquid nitrogen and homogenizing in 5 ml g–1 of extraction buffer for 1 h at 4°C (50mM Tris-HCl pH 7.5, 150mM NaCl, 10% Glycerol, 0.2% NP-40, 5mM MgCl2, 5mM DTT, containing one tablet per 10 ml of protease inhibitor cocktail, Roche). Cell debris was removed by centrifugation at 12 Krpm at 4°C for 20 min and the supernatant was recovered. Protein extract was pre-cleared by incubation with 10 μl Protein A Dynabeads (Invitrogen) at 4°C for 30 min. Supernatant was recovered by centrifugation at 12 Krpm at 4°C for 1min. Precleared extracts were incubated with AGO9 antibody (1:100 dilution) and 30 μl of Protein A Dynabeads at 4C overnight. The immunoprecipitate was pelleted and washed three times (15 min each in rotator shaker) in extraction buffer. Small RNAs from the purified AGO9 complex were isolated using Ambion columns. Small RNA was resolved on a 12.5% denaturing PAGE 7M-Urea gel and stained with SYBR-gold (Invitrogen). For cloning, gel slices within the range of 18–30 nt were excised, and the RNAs were eluted and purified as described above. Small RNAs were sequenced by Fasteris (Switzerland).

### Flow cytometry and staining of Arabidopsis pollen

Flow cytometry and Alexander staining was performed as previously reported^65^. Pollen was collected in 1.5mL eppendorf tubes by vortexing open flowers in pollen extraction buffer (PEB, 10 mM CaCl2, 2 mM MES, 1 mM KCl, 1% H3BO3, 10%) for 3 min 47, followed by filtration through a 30um mesh (Partec/Sysmex) and centrifugation at 5,000g for 1 min. Pollen was suspended in 50ul of PEB, immediately frozen in liquid nitrogen and stored at -80C until RNA extraction. Open flowers were collected into a 50 ml falcon tube, frozen in liquid nitrogen, and stored at -80C for storage. The pollen enriched fraction was isolated with the procedure described in Borges, et al. ^65^ using Sperm Extraction Buffer (SEB). Intact pollen grains were sorted with fluorescence-activated cell sorting (FACS) using a S3e Cell Sorter Bio-Rad. To the obtained pollen sample in 1.5 ml tube, 100 μl of acid-washed glass beads (425–600 μm, Sigma) was added and vortexed continuously at medium speed for 1 minute in order to break mature pollen grains. The sample was transferred to a new tube and the volume of the sample was made up to 1 ml with SEB. 1 µl of Sybr Green was added for staining. The sample was ready for FACS for sperm cells sorting according to Borges, et al. ^65^. RNA from the sperm cells fraction was isolated with Trizol, treated with DNase I, and used for small RNA library preparation (NextFlex Small RNA Kit, Perkin Elmer). Pollen staining was done following Alexander’s staining protocol^66^. Only viable pollen grains take up the stain.

### Bioinformatic analysis of rRNA coverage

Adapters were trimmed from reads, converted to fasta and collapsed using fastx toolkit. Trimmed reads between 15-30nt in length were mapped to the Arabidopsis genome (TAIR10) using bowtie^67^ allowing mismatches and multi-mapping. Coverage across the rRNA locus on chromosome 2 (position 3706-9521) was calculated using bedtools^68^. Coverage within a sample was normalized to the total amount of coverage across the locus, and the log2 fold-change between paired samples (IP/Unbound, CMC/Mock) was calculated and averaged between technical replicates. Sites of predicted pseudouridylation were taken from published datasets and the average log2 fold change surrounding these sites was calculated^69,70^.

### Bioinformatic analysis of CMC/Mn2+-treated Arabidopsis samples

For adapter trimming and size selection of reads (18-42 nt) Cutadapt was used^71^. Terminal deletions in CMC+ samples were detected using DeAnnIso^72^ or miRMOD^73^. For internal deletions detection reads were aligned to the Arabidopsis genome (TAIR10) using BWA^74^ and further processed using SAMtools^75^ bam-readcount and AWK homemade scripts. For identification of Ψ-modified RNA bases in the CMC/Mn+ protocol the following criteria were used: at least 5 reads for a particular miRNA and at least 1.5 CMC+/CMC-fold change for the ratio between read with mismatch/all reads for a particular miRNA.

### Bioinformatic analysis of Arabidopsis siRNA/miRNA

Adapters were trimmed from reads, converted to fasta and collapsed using fastx toolkit. Trimmed reads between 20-25nt in length were mapped to the Arabidopsis genome (TAIR10) using bowtie allowing mismatches and discarding unmapped reads. Individual sequences with an average of >10 reads per sample were selected and analyzed using DESeq2^61^, treating CMC+/Mock or IP/Unbound fractions as pairs. Mature miRNA sequences from miRBase (v21) were extracted from these datasets. Sequences mapping to tRNA, snoRNA, snRNA or rRNA were removed for analysis of 5′ and 3′ bias. UNUAR-containing and 3ʹCCA tRFs were identified by extracting reads mapping to tRNAs and selecting those containing the UNUAR motif or 3ʹCCA; these sequences were extracted from the DESeq2 output for plotting.

### Bioinformatic analysis of mouse miRNA

The mouse small RNA Sequencing data from NIH/3T3 cells were analyzed with the SeqImp pipeline configured with the mm10 assembly of the Mus musculus genome, the transcriptome annotation from Ensembl (v89) and the miRbase annotation of microRNAs (v21). The pipeline was executed with the default mouse miRNA configuration and default parameters, specifying the sequence of the 3′ adapter used for library preparation (AGATCGGAAGAGCACACGTC). The miRNA quantifications obtained with SeqImp were further analyzed in R/Bioconductor with the DESeq2 package^61^. Briefly, counts were library-size normalized using the median ratio method as implemented in the estimateSizeFactors function. Variance was estimated with the estimateDispersions function and statistical testing for enrichment performed with the Wald test as implemented in the nbinomWaldTest function. P-values were corrected for multiple hypothesis testing with the Benjamini-Hochberg procedure.

### Analysis of TE-derived siRNA

Collapsed reads 18-30 nt long were mapped to the TAIR10 genome allowing multimapping, but no mismatches using Bowtie. Quantification of siRNA mapping to each TE was performed as described previously^9^. Reads mapping to TE families were aggregated and the average log2 fold-change between paired samples was calculated. The mean of all technical replicates (generated by separate CMC/mock or IP/unbound library preparations on the same RNA) for any given biological replicate was taken prior to calculating the mean of all biological replicates for each tissue (inflorescence/pollen) and treatment (CMC or IP) combination.

### Metaplots

Uncollapsed reads 20-25nt in length were aligned to the Arabidopsis (TAIR10) genome using Bowtie allowing mismatches but only a single mapping event per read. Mapping files from the same treatment (i.e. CMC+, CMC-, IP, Unbound) were merged after normalization (as above) to enable the production of a single representative plot. Coverage was calculated using bedtools^68^. Metaplots were produced using the Versatile Aggregate Profiler^76^ using a window size of 25, and a smoothing window of 6 for TE analysis, and a window size of 50 and smoothing window of 12 for DME target analysis. For comparison between psd-13 and wild-type, metaplots of three biological replicates were created and the mean enrichment was plotted.

### Bioinformatic analysis of mouse testis small RNA

Reads were clipped for Illumina Truseq (AGATCGGAAGAGCACACGTCTGAAC) or Perkin Elmer Nextflex Small RNA adapter sequences (N(4)TGGAATTCTCGGGTGCCAAGG) using Cutadapt. Reads were required to be 14-44nt and have a quality score of at least Q20 over 90% of their length. tRNA fragment sequences (tRFs) were analyzed as described^77^ using all tRNA sequences included in the current UCSC repeat annotation (ftp://hgdownload.soe.ucsc.edu/goldenPath/mm10/database/rmskOutCurrent.txt.gz). Reads that were not tRFs were mapped to the UCSC mm10 genome sequence using Bowtie 2, filtered for 0-3 mismatches (to accommodate mismatches due to RNA modifications) using SAMtools and BAMtools, and intersected with the UCSC repeat annotation using BEDtools. Repeat and non-repeat reads were further intersected with mm10 mouse piRNA clusters (https://www.smallrnagroup.uni-mainz.de/piCdb/) to result in piRNA reads that overlap (i) transposon sequences, (ii) structural RNAs or (iii) neither. Raw counts of individual piRNA sequences were analyzed for pseudouridine enrichment (enriched in IP, depleted in CMC-treated libraries) using DESeq2^61^. Similarly, raw counts per piRNA cluster were analyzed using DESeq2, but each cluster was required to have at least an average of 100 raw reads (per IP or per CMC-untreated sample). Read counts that were used as input for DESeq2 analysis were from three replicates of each condition (input, IP; CMC-untreated, CMC-treated). Log2 fold changes were plotted against -log10(adjusted p-value) using R.

## Acknowledgments

The authors would like to thank J.S. Parent and M. Donoghue for discussions and observations, Joe Simorowski, Uma Ramu and Alex Oruci for technical assistance, and Tim Mulligan for plant care.

## Author contributions

R.P.H., J.D., C.S.A., V.M., T.K. and R.A.M. conceived and designed experiments. R.P.H., V.M., J.D., C.S.A., A.L, F.V., and A.H. performed the experiments. R.P.H., J.D., T.L. and A.J.S. performed bioinformatic analysis. F.B. provided code and advice for transposon analysis. R.A.M. and R.P.H. wrote the manuscript with input from V.M., J.D., C.S.A., A.J.S., T.L. and T.K.

## Data and materials availability

Data has been deposited in GEO under accession numbers GSE229986 (reviewer token gbetuskozpalzgh); GSE230359 (Reviewer token ufarcgsafjczjgh); GSE230228 (Reviewer token cnmnamgefjyjxmt); GSE222751 (Reviewer token cjcjommurpohpav).

## Supplementary Materials

Supplemental Table S1

Supplemental Data S1 to S10

## Extended Data

**Extended Data Figure 1.**
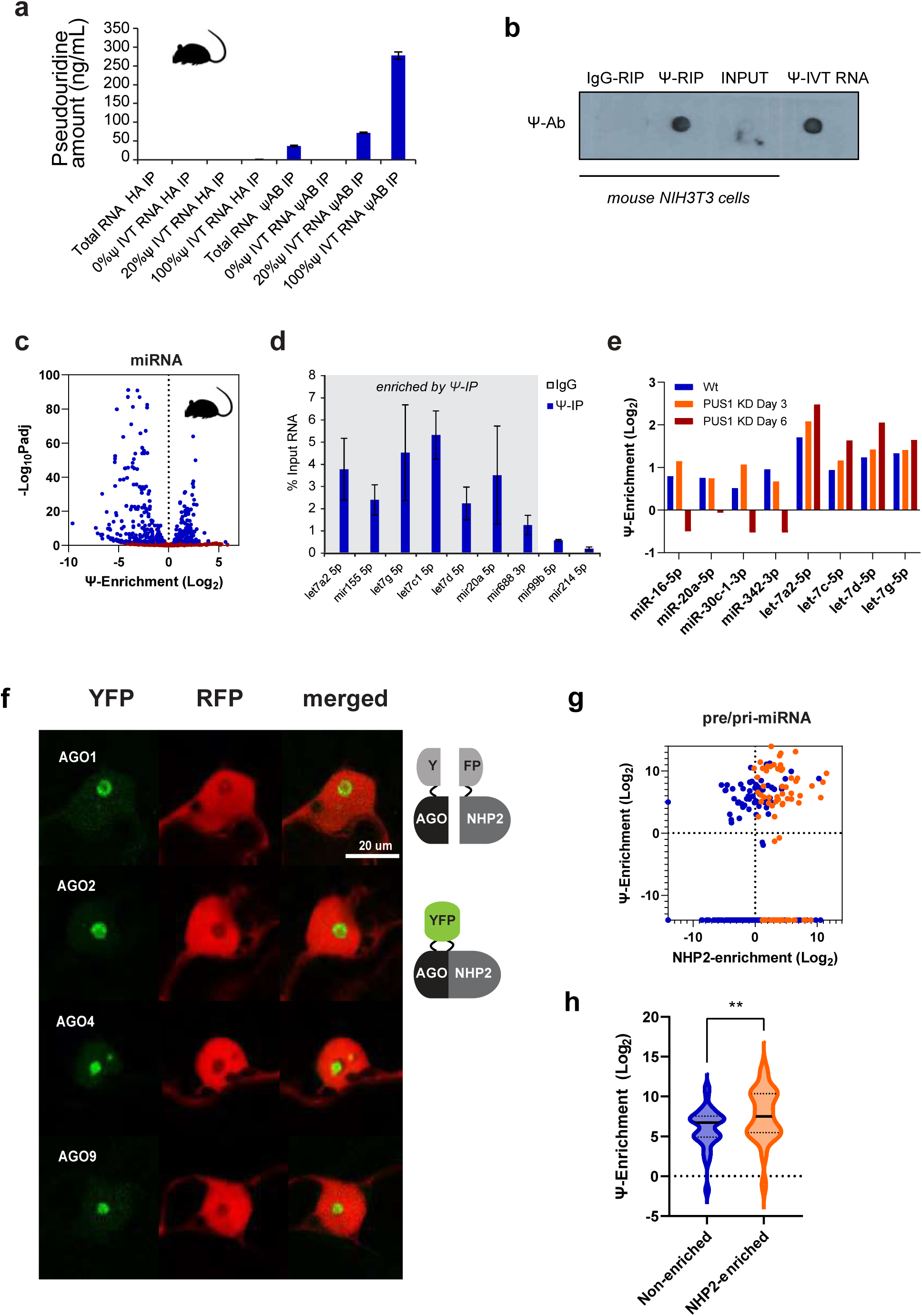
miRNA from plants and mammals are pseudouridylated. (**a**) Immunoprecipitation with anti-Ψ antibody enriches for Ψ-containing RNAs produced through in vitro transcription (IVT) as determined by mass spectrometry (n = 3 biological replicates, error bars ± SD). (**b**) Immuno-dot blot of 100ng total input RNA or 100ng RNA immunoprecipitated with anti-Ψ antibody or control immunoglobulin G (IgG). 100ng of 100% Ψ IVT RNA was used as control. (**c**) Volcano plot of miRNAs enriched by small RNA immunoprecipitation using a Ψ-specific antibody (Ψ-IP) in mouse fibroblast NIH/3T3 cells compared to input fractions. Significantly enriched (p<0.01) miRNAs are highlighted in blue. (**d**) Validation of enriched miRNAs using qRT-PCR. (**e**) Ψ-enrichment of miRNAs affected by PUS1 knockdown (KD) 3 days and 6 days after PUS1 shRNA transfection. (**f**) AGO proteins and Dyskerin subunit NHP2 were fused at the N-terminus to N- and C-terminally split YFP, respectively. Positive interaction is shown by YFP signal, with 35S::RFP as an expression control. (**g**) Ψ-enrichment of precursor pri-miRNAs in Arabidopsis flower buds plotted against pri-miRNAs bound to NHP2 (blue) and detected by RIP-seq in 2 or more replicates (orange). (**h**) Violin plot showing significant difference between NHP2-enriched and non-enriched pri-miRNA and Ψ enrichment of the mature miRNA by Ψ -IP (Log_2_ IP/IgG; (one-way ANOVA; **p<0.01).

**Extended Data Figure 2.**
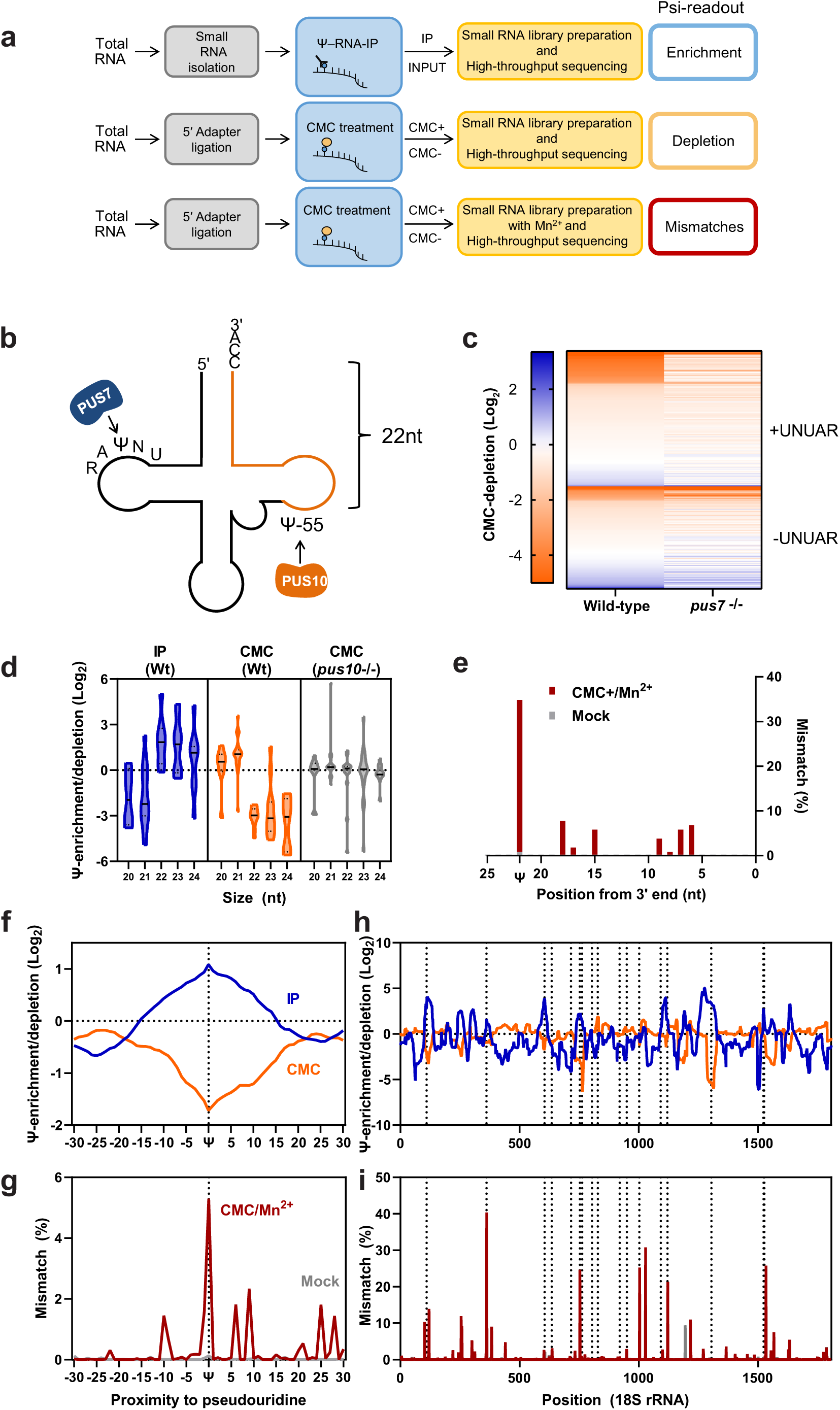
Strategies for robust detection of Ψs in small RNAs. (**a**) Three strategies to detect Ψ in small RNAs and their predicted effect on pseudouridylated sequences. Details are in Methods (**b**) Known sites of pseudouridylation in tRNAs mediated by PUS7 (targets UNUAR) and PUS10 (targets Ψ-55). Ψ-55 is 22 nt “upstream” of the 3’ end of -CCA tailed tRNA fragments. (**c**) Heatmap showing individual tRNA fragments (tRFs) with and without the UNUAR motif in wild-type and *pus7*-/- libraries treated with CMC. (**d**) 3′CCA tRFs ≥22 nt contain canonical Ψ-55 and are enriched by anti-Ψ antibody and depleted by CMC treatment, whereas shorter fragments do not contain this nucleoside and are not enriched/depleted. Ψ-55 is ablated in mutants of PUS10. (**e**) Deletions in 3ʹ CCA tRFs based on position in CMC/Mn^2+^- and mock-treated libraries. (**f, g)** Metaplots showing coverage depletion/enrichment in CMC/IP libraries at rRNA pseudouridylation sites (f) compared to mismatches from Mn2+/CMC+ libraries (g). Average across 42 predicted pseudouridylated sites in 18S and 25S rRNA and from 17 predicted sites in 18S rRNA (CMC mismatch/Mock mismatch). (**h, i**) Enrichment/depletion/mismatches across 18S rRNA in CMC/IP (h) and CMC/Mn^2+^ (i) libraries. Predicted Ψs are marked with dotted lines.

**Extended Data Figure 3.**
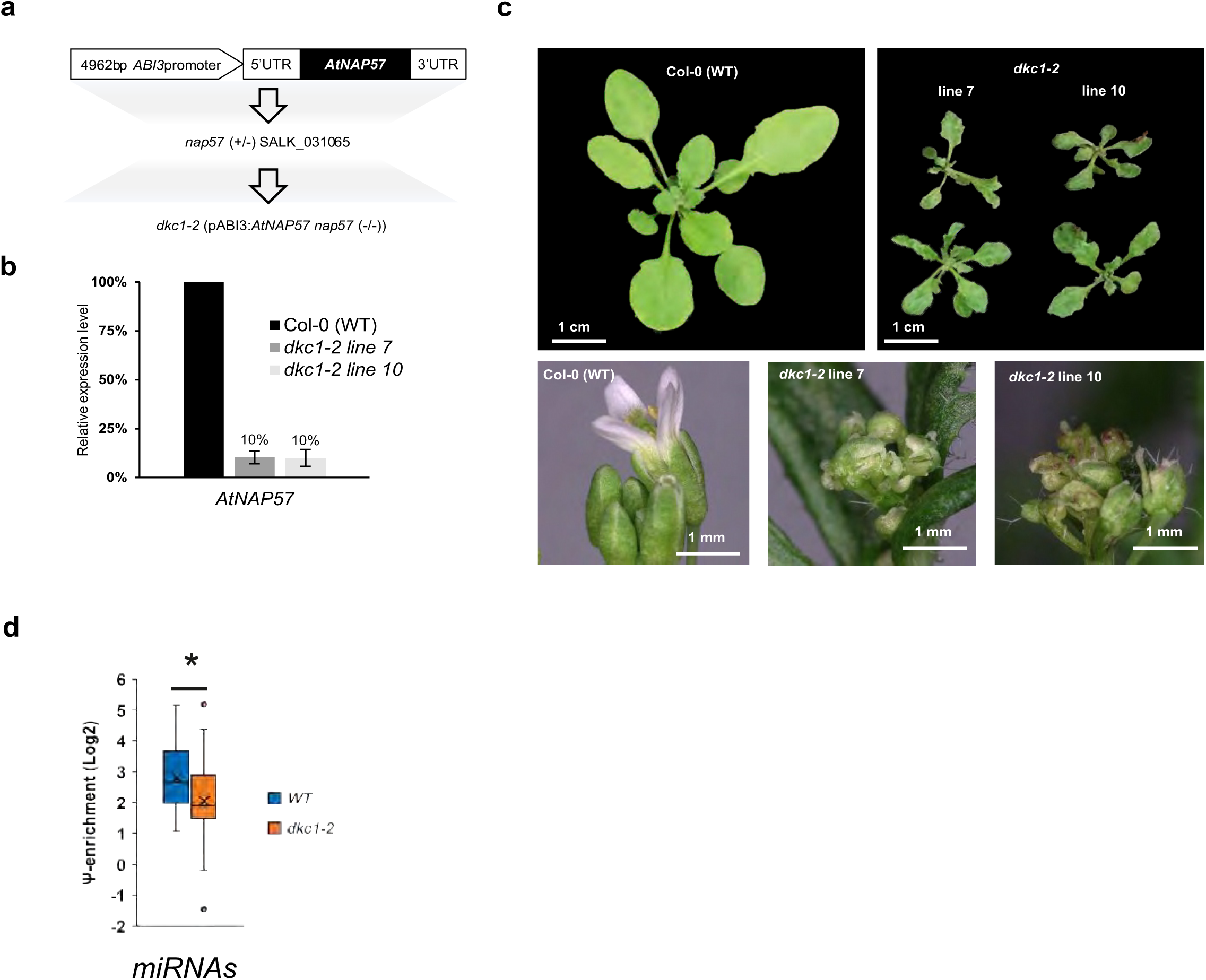
Arabidopsis mutants in Dyskerin/NAP57. (**a**) Generation of *dkc1-2* transgenic plants*. nap57* (SALK_031065) was complemented with the pABI3:AtNAP57 transgene (**b**) AtNAP57 transcript level in *dkc1-2* transgenic lines analyzed by qPCR (n=3) (**c**) *dkc1-2* transgenic plants show severe developmental defects including dwarfism, abnormal leaf shape, deformed and infertile flowers. (**d**) Enrichment of miRNAs by Ψ-IP in wild-type and *dkc1-3* leaves based on DESeq2 analysis.

**Extended Data Fig. 4.**
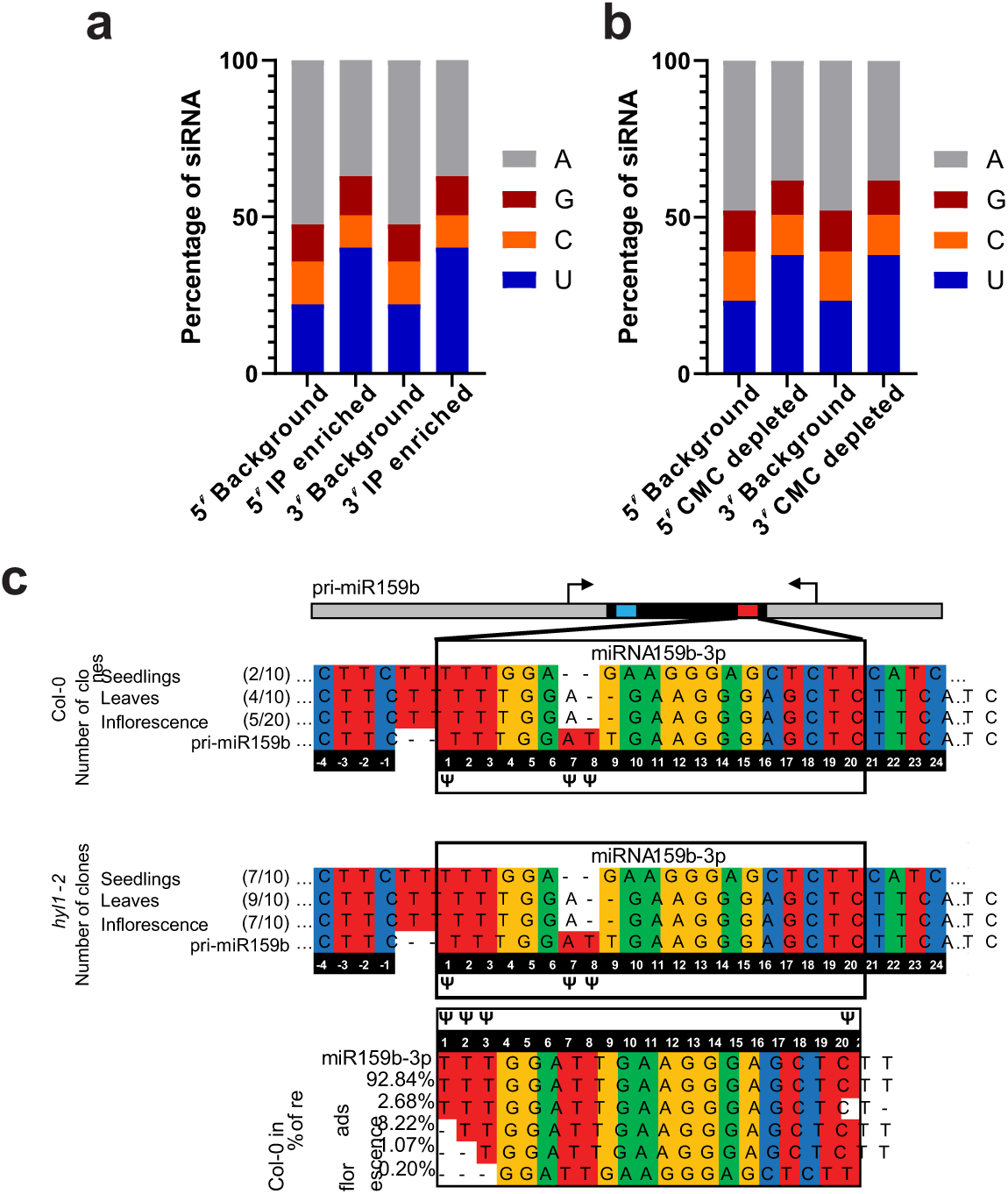
Mapping of miRNA pseudouridylation sites in Arabidopsis by CMC labeling and reverse transcription in the presence of Mn^2+^ (CMC/Mn^2+^). (**a**, **b)** Bar chart showing proportions of 5ʹ and 3ʹ nucleotides in (a) Ψ-IP-enriched and (b) CMC-depleted siRNAs from inflorescence compared to background (non-structural) small RNAs. IP-enrichment and CMC-depletion show biases towards 5ʹ and 3ʹ U. (**c**) Sites of predicted Ψ (insertions and deletions) in precursor pri-miR159b PCR amplicons (upper panels) and mature miR159b (lower panel; high-throughput sequencing) in seedlings, leaves and flower buds of wild-type and *hyl1-2* microprocessor mutants. pri-miRNA amplicons (upper) and mature small RNA sequences (lower) had insertions and deletions at predicted sites. Number of clones with the respective amplicon is shown in brackets.

**Extended Data Figure 5.**
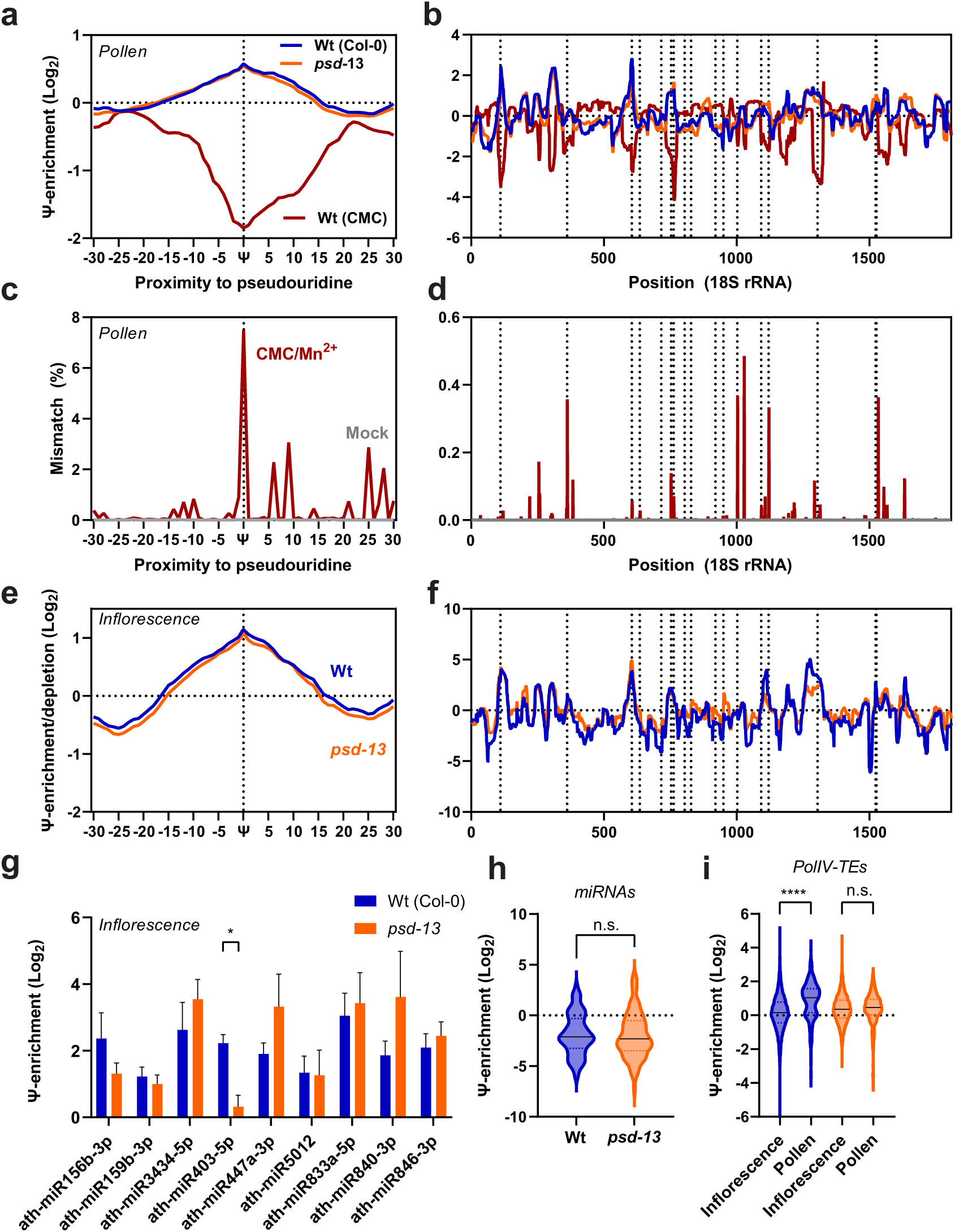
Pseudouridine in miRNAs and TE-derived siRNAs in pollen from WT and *paused (psd-13)/*exportin-t mutants. (**a**) Metaplots of read coverage at 42 known pseudouridylated sites in 18S and 25S rRNA, relative to untreated controls, in CMC-treated WT (magenta) and Ψ-IP WT (blue) and *psd-13* (orange) small RNA sequencing libraries from pollen. (**b**) Read coverage from the same libraries at 17 rRNA pseudouridylation sites in 18S rRNA. (**c, d**) metaplot and browser analysis of predicted sites in rRNA from CMC/Mn^2+^ analysis relative to mock treated controls. (**e, f**) Metaplots (e) and read coverage (f) relative to untreated controls in WT (blue) and *psd-13* (orange) Ψ-IP small RNA sequencing libraries from inflorescence. Known positions of Ψ in b, d and f are marked with dotted lines. (**g**) Ψ-enrichment of individual miRNA detected by Ψ-IP and small RNA sequencing in WT and *psd* mutant inflorescence (n=2 biological replicates). (**h**) Ψ-enrichment of miRNA overall in WT and *psd* mutant inflorescence (one-way ANOVA, p>0.05) (**i**) Violin plot of Ψ-enrichment of siRNA from individual TEs based on IP-enrichment (Log2 IP/Unbound) in flower buds (inflorescence) and pollen from wild-type and *psd* mutants. Significant Ψ enrichment was observed in pollen, but not in inflorescence, and depended on *PSD* (p<0.001, one-way ANOVA).

**Extended data Figure 6.**
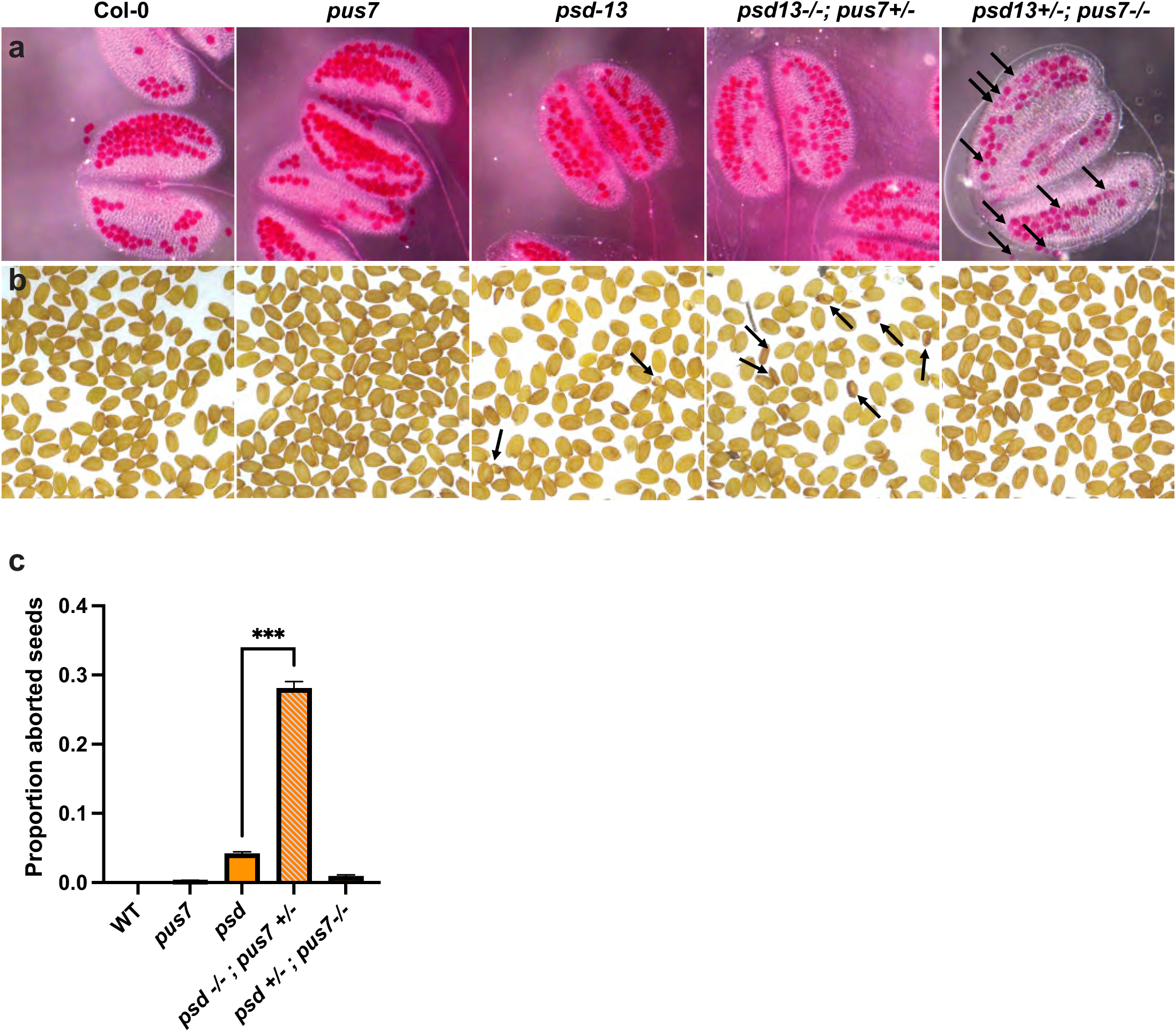
Mutation of *psd* and *pus7* is synthetically lethal in Arabidopsis. (**a**) Alexander staining of pollen from Col-0, *pus7*, *psd-13*, *psd-13*-/-; *pus7*+/-, and *psd-13*+/-; *pus7*-/- parents. *psd*-13+/; *pus7*-/- had semisterile pollen abortion (loss of Alexander staining, arrows). (**b**) Mature seeds from each parent. Aborted seeds in *psd-13*are highlighted with arrows. (**c**) Quantification of approximately 500 seeds from each genotype revealed *psd*-13-/-; *pus7*+/- have 25% seed abortion, significantly more than *psd-13* alone (<1%; p<0.001). Double homozygous mutants could not be obtained (n>150 plants).

**Extended Data Figure 7.**
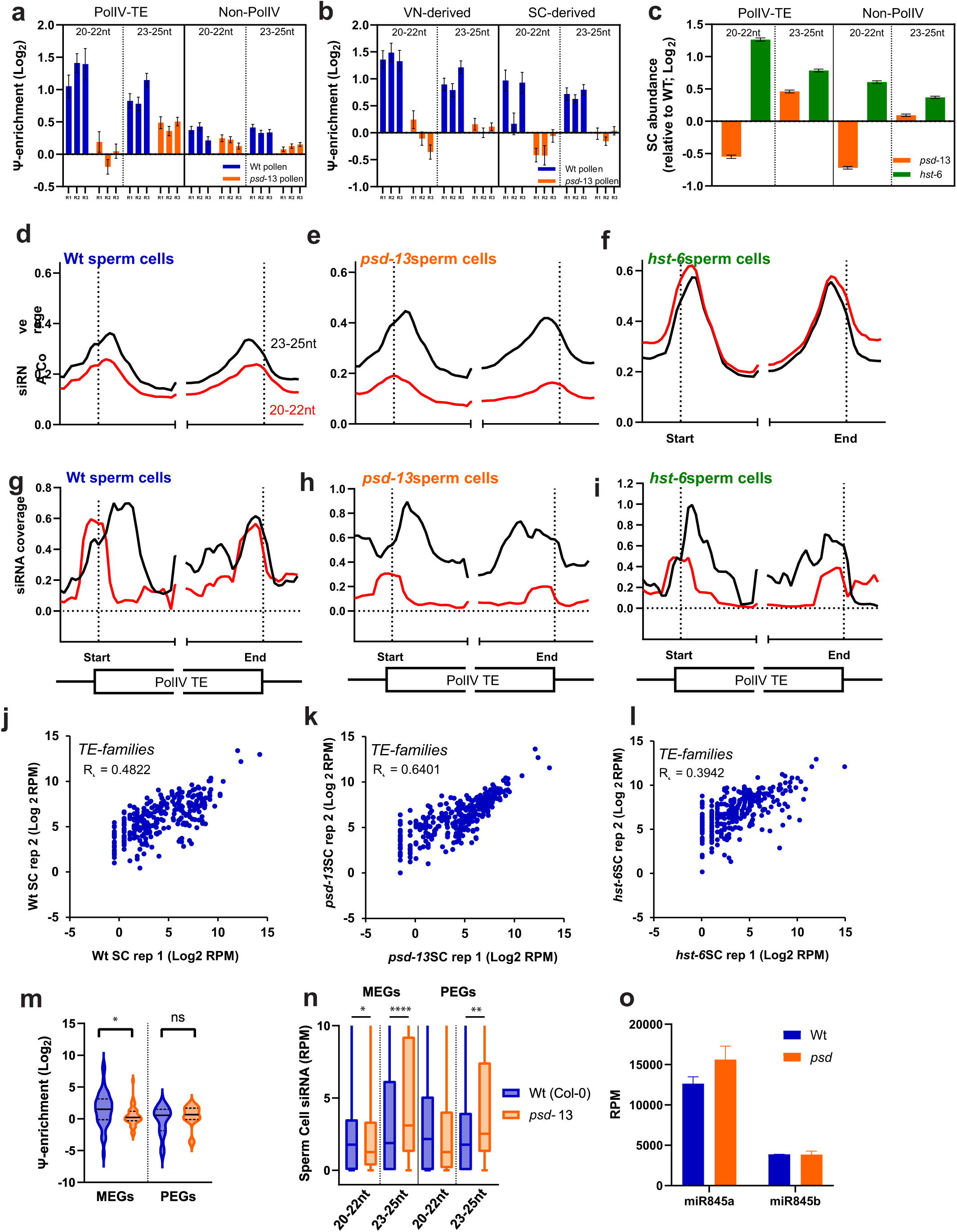
Characterization of pseudouridylation of siRNA in *psd-13* mutants. (**a,b**) Ψ-enrichment of PolIV-dependent and -independent (a) or VN- and SC-derived (b) TE-siRNA in wild-type and *psd*-13 pollen for each size class (average enrichment of TEs from each classification is shown +/- SEM for each biological replicate). (**c**) siRNA abundance was determined by sequencing small RNA from FACS sorted wild-type, *psd*-13 and *hst*-6 sperm cells. siRNA derived from PolIV-dependent and non-PolIV TEs was assessed in each size class (average enrichment of TEs relative to wild-type from each classification is shown +/- SEM). (**d-i**) Metaplots of 20-22nt (red) and 23-25nt (black) siRNA abundance across PolIV-TEs from biological replicates of wild-type (a, d), *psd-13* (b, e) and *hst-6* (c, f) sperm cells sorted by FACS. 250 bp up-and down-stream regions as well as 500 bp from each end of the TE are shown. (**j-l**) Scatterplots of small RNA sequencing data from two biological replicates mapped to TE families in (j) wild-type, (k) *psd-13* and (l) *hst-6* sperm cells isolated by FACS. (**m**) Ψ-enrichment of siRNAs from TEs surrounding maternally expressed imprinted genes (MEGs) and paternally expressed imprinted genes (PEGs) in wild type (blue) and *psd-13* (orange) pollen. * = p<0.05, ANOVA. (**n**) Abundance of 20-22 nt and 23-25 nt siRNA surrounding imprinted genes in wild-type (blue) and *psd-13* (orange) sperm cells. siRNA in sperm cells matching MEGs and PEGs was lower in *psd-13* mutants for 20-22nt siRNA but relatively higher for 23-25nt siRNA. (**o**) levels of miR845a and miR845b were unchanged or higher in *psd-13* mutant pollen (error bars show SD of biological replicates, n=3)

**Extended Data Fig. 8.**
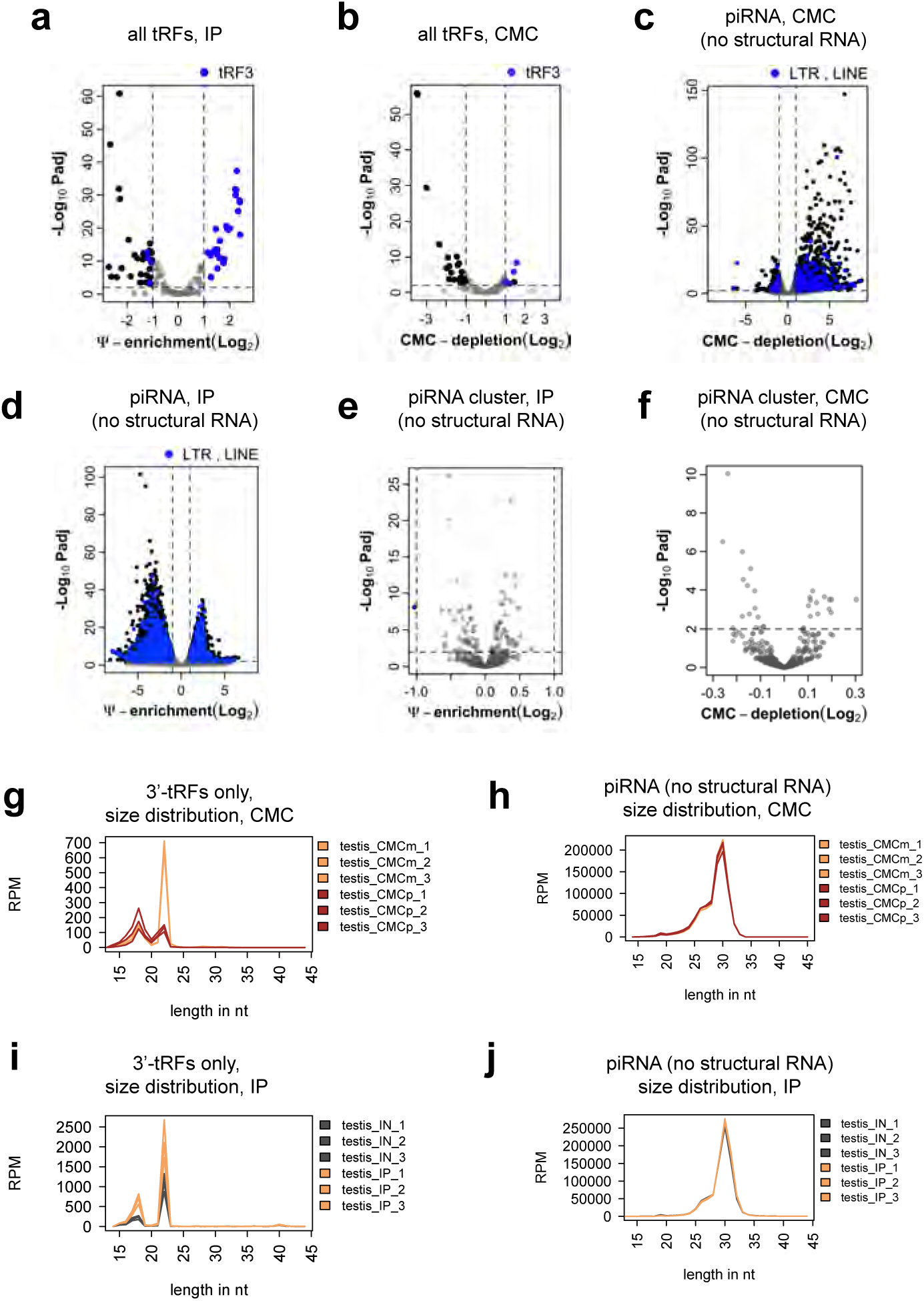
Pseudouridine in piRNAs and tRNA fragments (tRFs) from mouse testis. **(a, b)** Ψ enrichment by (a) Ψ-IP and (b) CMC-depletion of small RNA from testis from 8 weeks-old male mice. Each dot corresponds to tRFs from a single tRNA anticodon isotype. Statistically significant (p-adj. <0.01, log2 fold change > |1|) enriched/depleted tRFs are in black, 3’-tRFs are colored blue; grey dots mark tRFs without significant enrichment/depletion. **(c)** Volcano plot of individual piRNA sequences in testis (8 weeks old males) by CMC-depletion and sequencing. Black: significant fold changes (p-adj. <0.01, log2 fold change > |1|); grey: not significant; blue: significantly enriched/depleted piRNAs overlapping LTR and LINE transposon sequences. (**d**) Volcano plot of individual piRNA sequences. Black: significant fold changes (p-adj. <0.01, log2 fold change > |1|); grey: not significant; blue: significantly enriched/depleted piRNAs overlapping LTR and LINE transposon sequences. (**e,f**) Volcano plot of read counts per piRNA cluster using (e) Ψ-IP and (f) CMC-detection. **(g,h)** Size distribution of (g) 3’-tRFs and (h) piRNA sequences (without structural RNAs) using CMC-depletion and sequencing. **(i, j)** Size distribution of (i) 3’-tRFs and (j) piRNA sequences (without structural RNAs) using Ψ-IP and sequencing.

